# Neural variability compresses with increasing belief precision during Bayesian inference

**DOI:** 10.1101/2024.01.11.575180

**Authors:** Alexander Skowron, Julian Q. Kosciessa, Robert Lorenz, Ralph Hertwig, Wouter van den Bos, Douglas D. Garrett

**Affiliations:** Max Planck UCL Centre for Computational Psychiatry and Ageing Research, Berlin/London; Center for Lifespan Psychology, Max Planck Institute for Human Development, Berlin, Germany; Humboldt Universität zu Berlin, Germany; Donders Institute for Brain, Cognition and Behaviour, Nijmegen, The Netherlands; Lise Meitner Group for Environmental Neuroscience, Max Planck Institute for Human Development, Berlin, Germany; Center for Adaptive Rationality, Max Planck Institute for Human Development, Berlin, Germany; University of Amsterdam, The Netherlands

## Abstract

To make optimal decisions, intelligent agents must learn latent environmental states from discrete observations. Bayesian frameworks argue that integration of evidence over time allows us to refine our state belief by reducing uncertainty about alternate possibilities. How is this increasing belief precision during learning reflected in the brain? We propose that moment-to-moment neural variability should scale with the degree of reduction of uncertainty during learning. In a sample of 47 healthy adults, we found that BOLD signal variability (SD_BOLD_, as measured with functional MRI) indeed compressed with successive exposure to decision-related evidence. Crucially, more accurate participants expressed greater SD_BOLD_ compression primarily in Default Mode Network regions, possibly reflecting the increasing precision of their latent state belief during more efficient learning. Further, computational modeling of behavior suggested that more accurate subjects held a more unbiased (flatter) prior belief over possible states that allowed for larger uncertainty reduction during learning, which was directly reflected in SD_BOLD_ changes. Our results provide first evidence that moment-to-moment neural variability compresses with increasing belief precision during effective learning, proposing a flexible mechanism for how we come to learn the probabilistic nature of the world around us.

---

To make optimal decisions, animals often need to learn about “states” (i.e., decision-relevant properties) of the environment, such as the availability of fish in a nearby pond based on previous catch successes. These states are often not directly observable, so optimal agents need to combine the uncertainty about their beliefs with the evidence available to them. Research suggests that humans and other animals often take into account uncertainty during learning and decision-making^2-4^, while suboptimal use of uncertainty has been linked to maladaptive behavior observed in clinical and ageing populations^3^. Bayesian decision theory prescribes how agents should combine evidence with their internal beliefs under uncertainty to arrive at optimal estimates of external states and provides a benchmark to assess rational behavior in the context of learning ^2,3,5^. In this framework, an agent must represent a probability distribution over possible states. The variance of this distribution reflects the uncertainty of the agent’s state belief. As evidence accumulates and the agent refines its representation of the “correct” world state, the variance of the belief distribution is thought to reduce. How this increasing belief precision during learning is reflected in the brain remains unclear.

A potential candidate for tracking the degree of state uncertainty is the moment-to-moment variability of neural responses. At the level of sensory cortex, computational modeling and non-human animal work suggests that neuronal variability signals uncertainty about peripheral inputs, such that neural population activity reflects sampling from a probability distribution over potential stimulus features; the higher perceptual uncertainty, the higher the neural variability ^6-10^. This idea is in line with theoretic work suggesting that neural systems should maintain an appropriate degree of instability in the face of uncertainty to permit the exploration of alternative causes of incoming sensory input ^11^. While plausible, no study to date has investigated whether stimulus-evoked neural variability tracks changes in perceptual uncertainty in a learning task and how it relates to behavior. With regard to human data, a recent study by Kosciessa, et al. ^12^ showed that increasing uncertainty about the task-relevance of different perceptual stimulus features is accompanied by an increase in cortical “excitability” (heightened desynchronization of alpha rhythms and increased entropy in the EEG signal). However, it is not clear whether these perceptual accounts of uncertainty-based shifts in neural variability also translate to higher-order decision variables that arise while learning about latent environmental states.

In the context of learning, larger temporal variability under high uncertainty may not just represent exploration of potential world states but may allow more flexible updating of one’s belief once new information becomes available. Indeed, previous human work suggests that higher brain signal variability affords larger cognitive flexibility ^13,14^. For example, several studies have found that brain signal variability increases with increasing task demands (at least until processing limits are reached) and that the ability to upregulate variability predicts task performance ^15-22^. These studies commonly argue that neural variability supports performance under increasing task demand by allowing the brain to maintain flexible responding to stimulus information. In line with this idea, Armbruster-Genç, et al. ^16^ observed better performance on a task switching paradigm under higher BOLD signal variability, whereas cognitive stability during distractor inhibition was related to lower brain signal variability. During learning, task demands are highest early on, when the brain needs to maintain high flexibility to incorporate incoming evidence for belief updating. This should allow it to converge on the “correct” state representation with learning. We therefore hypothesize that moment-to-moment brain signal variability should compress with increasing belief precision (i.e., decreasing uncertainty) during learning (Figure 1).

**Figure 1.**
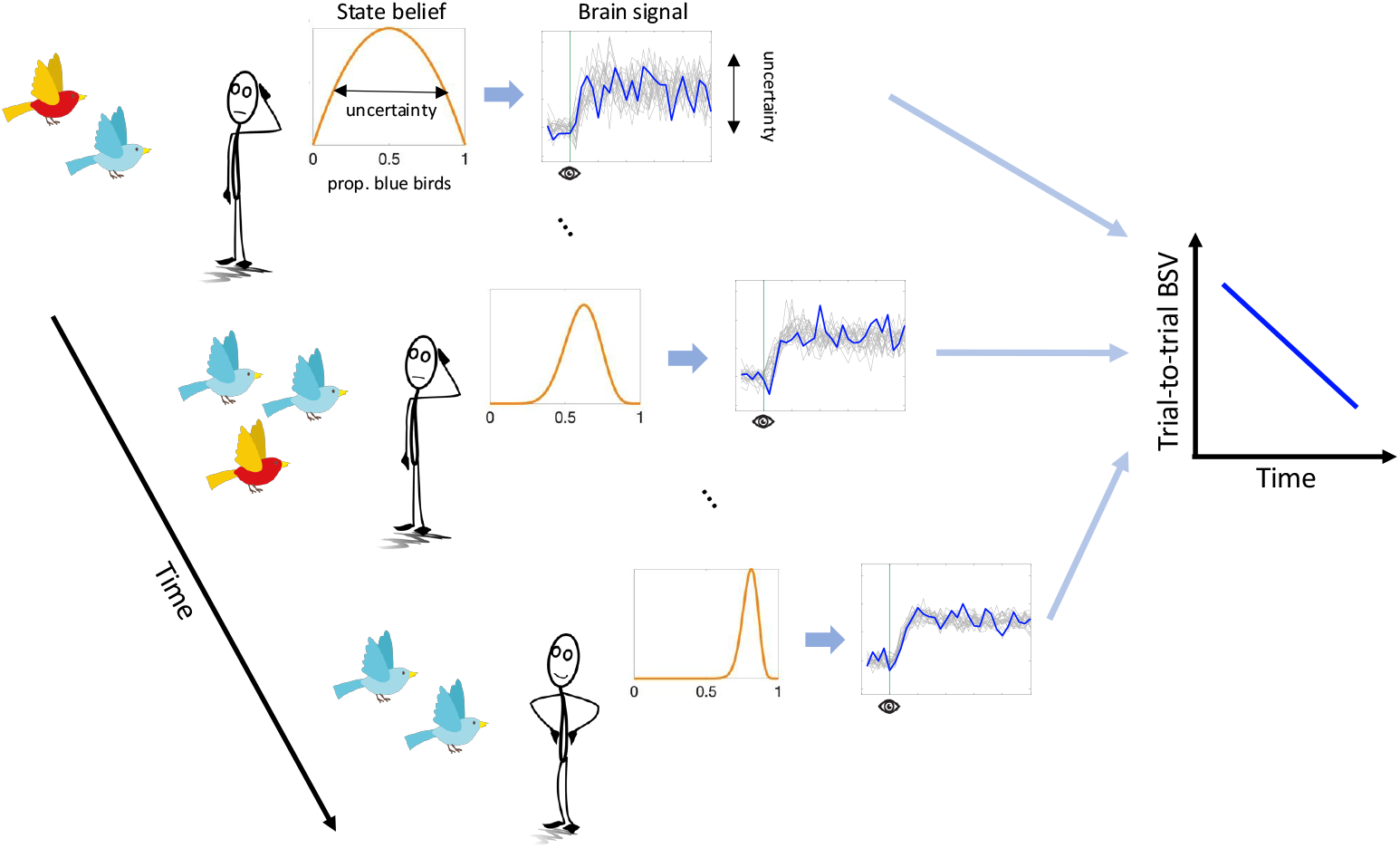
Hypothetical model of uncertainty and brain dynamics. In this example, the agent has to infer the proportion of blue birds in a world made up of red and blue birds by combining information across observations over time. Under high state uncertainty early on in the learning process, more moment-to-moment brain signal variability allows the brain to entertain a larger variety of possible states (i.e., proportions of blue birds in the world). With each new observation, the precision of the belief distribution increases and moment-to-moment brain signal variability compresses (plotted in blue). This hypothetical model also predicts more brain signal variability (BSV) across independent learning trials (plotted in gray) earlier than later in the learning process.

Notably, we assume that these neural dynamics unfold on a fast temporal scale, as suggested by electrophysiological evidence in human and non-human animals ^7,8,19-22^. However, within-trial dynamics should also affect neural variability across independent learning trials (see Figure 1): A more variable system should have a higher probability of being in a different state every time it is (sparsely) sampled. Conversely, when a system is in a less stochastic state, the within-trial variance is expected to reduce, yielding less across trial variance at the same time. This argument aligns with work by Orbán, et al. ^7^, who showed that a computational model of the sampling account of sensory uncertainty captures empirically observed across-trial variability of neural population responses in primary visual cortex. In the case of human research, this means that neuroimaging methods with slower sampling rates, such as functional MRI (fMRI), may be able to approximate within-trial neural variability from variability observed across trials. Indeed, the majority of previous fMRI studies reporting within-region, within-subject modulation of brain signal variability by task demand have exclusively employed block designs, necessitating that the main source of variability be between-rather than within-trial^15-18^.

In the current study, we acquired fMRI while participants performed a “marble task”. In this task, participants had to learn the probability of drawing a blue marble from an unseen jar (i.e., urn) based on five samples (i.e., draws from the urn with replacement). In a Bayesian inference framework, the jar marble ratio can be considered a latent state that participants must infer. We hypothesized that (1) trial-to-trial variability in the BOLD response (SD_BOLD_) would compress over the sampling period, thus mirroring the reduction in state uncertainty, and that (2) subjects with greater SD_BOLD_ compression would show smaller estimation errors of the jars’ marble ratios as an index of more efficient belief updating. A secondary aim of the current study was to directly compare the effect of uncertainty on SD_BOLD_ with a more standard General Linear Modelling (GLM) approach, which looks for correlations between average BOLD activity and uncertainty. This links our findings directly to previous investigations of neural uncertainty correlates, which disregarded the magnitude of BOLD variability ^2,23-37^. We hypothesized (3) that SD_BOLD_ would uniquely predict inference accuracy compared to these standard neural uncertainty correlates.

Our results showed that SD_BOLD_ closely tracked individual differences in the reduction of state uncertainty during learning which related to task accuracy in a unique manner that was not captured by a more standard GLM approach. As such, we identified a novel neural signal reflecting uncertainty during learning.

## METHODS

### Participants and procedure

51 young adults (24 female, 19-40 years [mean age = 26.37, SD = 3.74]) participated in the study. They were recruited from the participant database of the MPI for Human Development (Berlin, Germany) and gave written informed consent according to the guidelines of the German Psychological Society (DGPS) ethics board, who approved the study. All participants were right-handed, had normal or corrected-to-normal vision, did not concurrently participate in any clinical trials/ pharmacological studies, had no metal implants, did not take any centrally active medications, were not pregnant, and reported to be in good health with no known history of neurological, psychiatric or other severe (cancer, cardiac insufficiency, chronic liver/kidney disease, a severe hematopoietic condition, severe allergic/toxic reactions, brain/heart operation) health condition. Participants underwent fMRI while performing a perceptual gambling task. They received 10 Euros per hour in addition to a variable performance bonus of up to 2 Euros.

### The marble task

The task was divided into four blocks of 18 trials each. Participants performed three blocks during fMRI acquisition, while the final block was performed outside the scanner. We will only consider the three MR blocks for our analyses. Participants received instructions and completed nine practice trials prior to the first scanning block. Each trial of the task consisted of three phases: a sampling, an estimation, and a gambling phase. To investigate brain signal variability changes during learning, we focused on the sampling and estimation phases in this study (referred to as the “marble task”). A description of the gambling phase, which was unrelated to the learning process, is provided in the Supplementary Methods. The task design also included a between-trial reward manipulation. Here, we collapsed trials across the two levels of the reward manipulation given that we assumed those would only affect the choice and not the learning part. The task was programmed using Presentation software (Version 14.9, Neurobehavioral Systems Inc., Albany, CA, USA).

In the marble task, participants were asked to estimate the proportion of blue marbles in an unseen “jar” (i.e., urn), containing a total of 100 blue and red marbles, based on five successive samples (i.e., draws from the urn with replacement) presented successively during the *sampling phase* (Figure 2A). In total, 18 different jars with different proportions of blue marbles (ranging from 0.1 to 0.91) were presented across all trials (Supplementary table S1). Each jar was presented four times across the four experimental blocks in random order. Each sample from a given jar could contain either one, five or nine marbles, which manipulated the informativeness of the samples (i.e., one marble was least informative for inferring the proportion of blue marbles whereas nine marbles was most informative). Samples were presented in 3×3 grids with grey marbles serving as placeholders to ensure similar visual inputs across samples. Additionally, within each sample the order of the marbles (i.e., the location in the 3×3 grid) was permuted. The samples for each jar were randomly drawn from a binomial distribution with the corresponding probability of drawing a blue marble. This draw was performed once so each jar was associated with a consistent set of five samples with varying sizes (note that the total number of marbles across samples thus varied between jars). This ensured that all participants received identical sample information. For each trial with the same unseen jar, the order of the associated samples was varied randomly. Each sample was presented for 1s followed by a fixation cross presented for 2 to 6s. Following the sampling phase, participants were asked to indicate their estimate of the proportion of blue marbles in the jar by adjusting the ratio of blue to red marbles in a 100-marble grid using an MR-compatible five-button box. The starting value of the grid was always set to 50 red and 50 blue marbles. With the upper and lower buttons, the participants were able to adjust the grid in steps of five marbles, and with the left and right button in a one marble step, respectively. With the middle button participants confirmed their estimation. The maximum reaction time for the estimation phase was set to 7s. Participants received a bonus payment based on the average accuracy of their marble ratio estimates (performance-dependent bonus).

**Figure 2.**
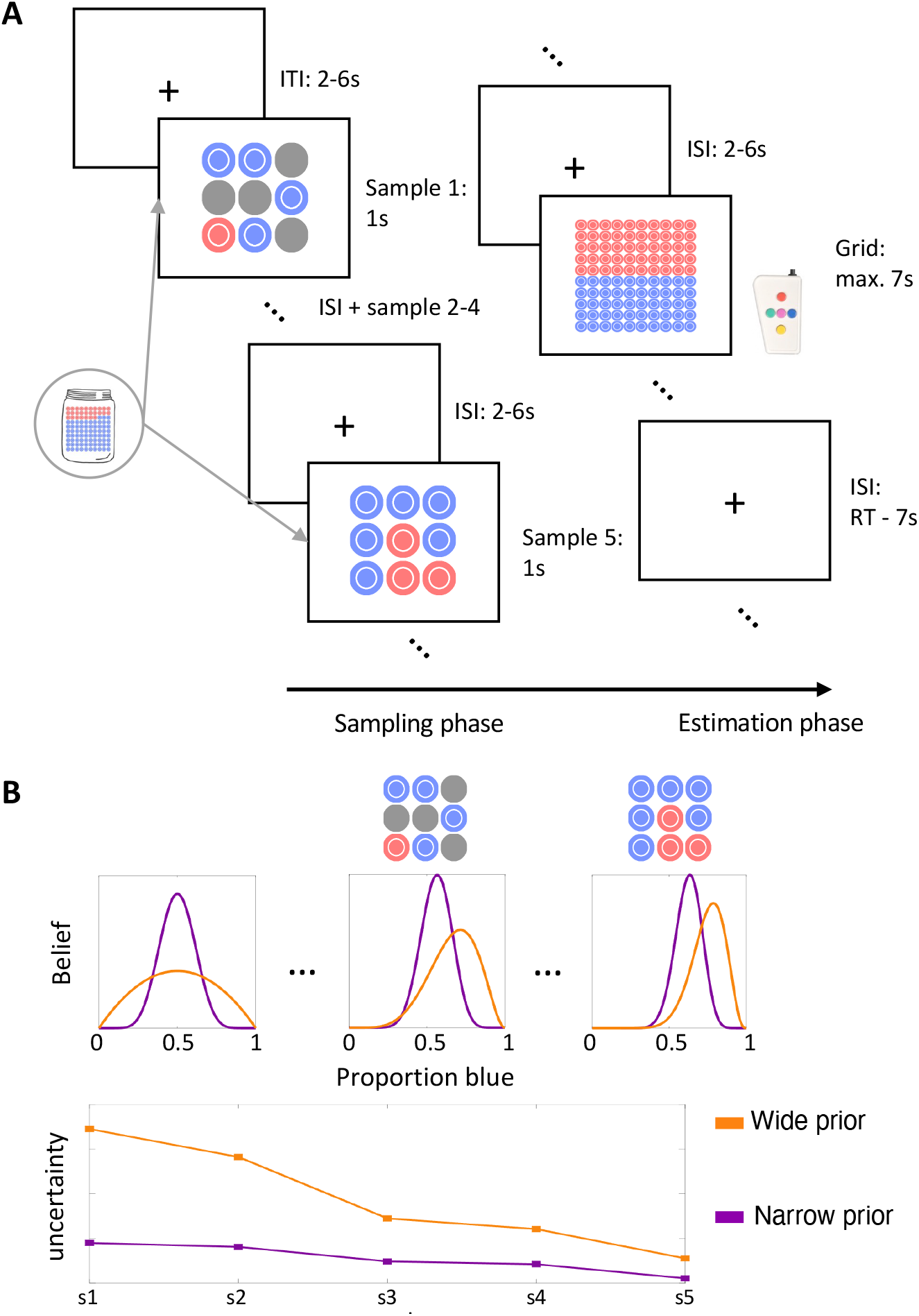
Behavior in the marble task. A: Task design. Participants observed five samples from an unseen jar (sampling phase) and then entered their estimate of the proportion of blue marbles in the jar into a 10×10 grid (estimation phase). B: Schematic illustration of the Bayesian observer model with varying priors. A narrower prior belief centered on a default 50:50 marble ratio predicts more misestimation of extreme jar proportions (top) and less uncertainty reduction during the sampling phase (bottom; s1 to s5 correspond to the five samples presented on a given trial).

### Behavioral modelling

We modeled participants’ responses using variants of a Bayesian observer model. The Bayesian observer represents the probability of drawing a blue marble from the unseen jar as a beta distribution with parameters *α* and *β*, which is updated after each draw according to Bayes’ rule (Figure 2B). Because the beta distribution is a conjugate prior for the binomial distribution (the likelihood function in this task), the posterior belief distribution is also a beta distribution with *α*_s+1_ = *α*_s_ + B_s_ and *β*_s+1_ = *β*_s_ + R_s_, where B_s_ and R_s_ are the number of blue and red marbles for a sample *s* respectively. The prior for the first draw of each trial is given by *α* = *β* = 1, representing a flat prior. The best estimate for the probability of drawing a blue marble is given by the expectation of the beta distribution 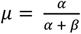.

To model empirical response patterns affecting subjects’ task accuracy, we considered two variants of this Bayesian observer model. In one model variant, we fit a parameter ρ ≥ 1 for each subject, which described the initial setting of *α* and *β* on each trial prior to observing samples from a given jar (where ρ = *α* = *β*; Figure 2B). A higher value for ρ describes a narrower prior around the default belief of 0.5. In a second Bayesian observer model variant, we fit a parameter *δ* ≥ 0, which exponentially weighs the evidence of each sample: *α*_s+1_ = *α*_s_ + (B_s_)^*δ*^ and *β*_s+1_ = *β*_s_ + (R_s_)^*δ*^. If *δ* < 1, the evidence of larger draws is underweighted, while for *δ* > 1 the evidence of larger draws is overweighted. Comparing these models allowed us to test whether deviations from the unbiased Bayesian observer model were related to prior representation or evidence weighting respectively.

To accommodate probabilistic responding of subjects, we compared these models with two alternative choice rules. One model family assumes that subjects’ reported estimates represent draws from the final beta distribution on each trial. Another model family assumes that subjects’ choices are draws from a truncated normal distribution (0 ≤ *x* ≤ 1) that is centered on the expectation of the final beta distributions: *N*(*E*[*beta*(*α, β*)], *σ*), where *σ* is a free model parameter that captures response noise around the model prediction. This parameter may reflect imprecision in entering one’s marble ratio estimate into the response grid, but it can also capture any unmodeled sources of (biased and random) errors in jar estimates beyond those explained by the model parameters of interest. This parameter may thus also capture model misfit.

Bayesian observer models assume that participants track the full belief distribution over potential marble ratios and consider uncertainty when updating their estimate. To test whether subjects indeed behave in a Bayesian manner, we also fitted a simpler Rescorla-Wagner model that only updates a point estimate of the marble ratio belief: 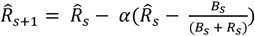, where 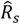 is the marble ratio point estimate after observing sample s, B and R_s_ are the number of blue and red marbles respectively, and 0 ≤ *α* ≤ 1 is the free learning rate parameter. To model subjects’ responses, we included a free noise parameter *σ* modelling Gaussian response noise around the final point estimate, similar to the noisy choice rule described previously for the Bayesian observer models.

#### Model fitting and comparison

We fitted behavioral models to participants’ responses on all trials of the three MR blocks by minimizing the joint negative log likelihoods under each model using the fmincon routine implemented in MATLAB (version 2017b). We ensured model convergence by fitting each model 10 times and used the fitting iteration with the minimal negative log likelihood across subjects. We compared model fits by computing the Bayesian Information Criterion (BIC): *BIC* = *k* ln(*n*) − 2 ln (*L*), where k is the number of parameters, n is the number of datapoints, and L is the maximized model likelihood. We report BICs for each subject and across all subjects.

#### Model simulations

We simulated behavior using the empirical parameter estimates and the trial sequences of each subject. This resulted in one synthetic dataset for each model with 51 simulated subjects each. We used these simulated datasets to perform model and parameter recovery checks. See the Supplementary Information for the results of these checks.

### Image acquisition and pre-processing

Participants underwent functional MRI scanning at the Max Planck institute for Human Development (Berlin, Germany) in a 3 Tesla Siemens TrioTim MRI system (Erlangen, Germany) using a multi-band EPI sequence (factor 4; TR = 645 ms; TE = 30 ms; flip angle 60°; FoV = 222 mm; voxel size 3 × 3 × 3 mm; 40 transverse slices). This amounted to 1010 T2*-weighted functional images per scanner run per subject (except for one subject, who had 1020 acquired images per run). A T1-weighted structural scan was also acquired (MPRAGE: TR = 2500 ms; TE = 4.77 ms; flip angle 7°; FoV = 256 mm; voxel size 1 × 1 × 1 mm; 192 sagittal slices).

T1-weighted images were brain extracted using ANTs software (version 2.3.5, http://stnava.github.io/ANTs/) using population level templates from the OASIS dataset (https://figshare.com/articles/dataset/ANTs_ANTsR_Brain_Templates/915436). The functional T2*-weighted scans were pre-processed in FSL (version 5.0.11 ^38,39^) FEAT separately for each run. The pipeline includes motion correction, brain extraction of the functional images, and spatial smoothing using a Gaussian 5mm kernel. Following FEAT, the functional images were first detrended using SPM’s detrend function (at a 3^rd^ order polynomial) and then high-pass filtered using a standard 8^th^ order butterworth filter implemented in MATLAB (version 2017b) with a cut-off of 0.01 Hz. We then performed ICA on the resulting data using FSL MELODIC ^40^ to identify residual artifacts. We manually labeled rejected components for 19 subjects (∼37% of the total data; for details on the component rejection criteria see ^41^) and then used these labels to train FSL’s ICA classifier FIX ^42,43^ to automatically label artifactual components for the remaining subjects. We tested different classification thresholds and validated classification accuracy for several randomly selected subjects from the test set. We accepted a threshold of 20 after this manual inspection. Rejected components were then regressed out of the data using FSL’s fsl_regfilt function. Lastly, functional images were first registered to the brain-masked T1-weighted images and then to MNI space (using the MNI152 template provided in FSL) via linear rigid-body transformation as implemented in FSL’s flirt function.

### Statistical analysis

#### Behavioral analysis

All behavioral analyses reported were run in SPSS (Version 25). To limit the effect of univariate outliers on the reported results, we winsorized the estimation error, extreme jar bias, and behavioral model parameter variables. Outliers were defined as datapoints that were 1.5 times the interquartile range beyond the 25^th^ or 75^th^ percentile respectively. We used the highest/lowest score of the non-outlying datapoints to impute outlying values. Between zero and three values were imputed for each variable.

#### Functional MRI analysis

Subject-level fMRI data were modelled using a mass-univariate GLM approach as implemented in SPM12 (Wellcome Department of Imaging Neuroscience, London, UK). To quantify BOLD response variability, we adapted a “least squares – single” (LS-S) approach as described by Mumford, et al. ^44^. This approach is implemented in an in-house version of variability toolbox for SPM (http://www.douglasdgarrett.com/#software) developed by our research group. This toolbox takes as input a standard GLM design matrix. We included the following regressors in each subject’s design matrix: Five regressors for the onsets of each successive sample presentation (with a duration of 1s), one regressor for the estimation phase onsets (with a duration of subjects’ RTs), and one regressor for the gambling phase onsets (with a duration of subjects’ RTs). All regressors were modelled as stick functions that were convolved with the canonical hemodynamic response function (HRF) and its first and second derivatives resulting in a total of 21 regressors per scanner run plus one constant regressors for each run. The toolbox then proceeds to iteratively fit GLMs which include one regressor modelling a single event and a second regressor modelling all other events of the same and all other conditions (this is done separately for each task run). To avoid issues with beta estimation close to the end of a timeseries (caused by truncation of the HRF), the toolbox discards any onsets that occur within the final 20s of the timeseries. Finally, the toolbox computes the standard deviation over the resulting (across-trial) beta estimates for each condition to yield the measure of SDBOLD used in this study. Compared to other approaches to SD_BOLD_ estimation, this SD_BOLD_ quantification allows one to parse dynamics amongst neighboring events/time points, while accounting for hemodynamic delays.

To compare our SD_BOLD_ results to standard analysis approaches, we also obtained restricted maximum likelihood (ReML) GLM beta estimates for each task condition. The design matrix of this GLM included the same regressors as the one passed to variability toolbox for SD_BOLD_ estimation. We defined contrasts to summarize condition effects across task runs. However, this approach ignores the variance in uncertainty trajectories across trials that result from our sample size manipulation (see Figure S1B). To account for this variance, we ran another GLM modelling the modulation of the BOLD signal by the posterior variance (i.e., uncertainty) of the unbiased Bayesian observer model during the sampling process. The design matrix of this GLM included the following regressors for each scanner run: one regressor for the sample onsets (with a duration of 1s), one regressor for the parametric modulation of the sample onsets regressor by model-derived posterior variances, one regressor for the estimation phase onsets (with a duration of subjects’ RTs), and one regressor for the gambling phase onsets (with a duration of subjects’ RTs). We mean-centered the parametric modulation regressor prior to model estimation. All regressors were modelled as stick functions (except for the parametric modulation regressor) that were convolved with the canonical hemodynamic response function (HRF) and its first and second derivatives resulting in a total of 12 regressors per scanner run. We also included three additional constant regressors modelling the mean of each scanner run. To look at the effect of the parametric modulation regressor, we defined a contrast that picked out the beta estimates of this regressor for each scanner run (more precisely, the parametric modulation regressors convolved with the canonical HRF).

To relate within-person SD_BOLD_ or standard GLM beta estimates to the task design or behavior, we use a partial least-squares approach (PLS) ^45^. PLS finds latent factors that express maximal covariance between brain and behavior/design matrices using singular value decomposition. Brain scores are defined as the product of the brain data matrix with their respective latent factor weights (saliences). We will refer to brain scores in our SD_BOLD_ analyses as “latent SD_BOLD_” and to the brain scores of our parametric modulation analyses as “latent uncertainty modulation”. To obtain a summary measure of the spatial expression of a latent variable, we can sum brain scores over all voxels. We performed 1000 permutation tests to assess the significance of the brain-design/behavior relationship against a null model. We then divided each voxel’s salience by its bootstrapped standard error, at +/-3 (approximating a 99.9% confidence interval), which serves as a pseudo-normalized measure of voxel robustness. We determined clusters of voxels with robust saliences by applying a cluster mask with a minimum threshold of 25 voxels. Prior to all PLS analyses, brain measures images were grey matter masked using the tissue prior provided in FSL at a probability threshold of > 0.37, and restrained to voxels with non-zero values for the respective brain measure across subjects.

To account for univariate outliers and non-linear relationships, we ran all reported brain-behavior analyses on ranked brain and behavioral variables in line with previous research ^46^. The results should thus be interpreted as monotonic rather than strictly linear relationships. Furthermore, we used a criterion of Cook’s distance > 4/N to identify multivariate outliers in all PLS models, removing where present. We nevertheless report the results of all analyses for the full dataset (N=51) without outliers removed in the Supplementary Results.

### Data availability

The code to perform all behavioral and neuroimaging analyses is available at: https://github.com/LNDG/Skowron_etal_2023. The research data required to replicate all reported results can be obtained by contacting the research data management unit at the Max Planck Institute for Human Development (Lentzeallee 94, 14195 Berlin, Germany) via this e-mail address. Interested parties will be required to provide a signed data handling agreement stipulating the conditions for data sharing before receiving access to the anonymized dataset.

## RESULTS

### Subjects’ estimation errors are explained by individual differences in prior belief width

Participants performed the “marble task” during functional MRI scanning (Figure 2A). On each trial, they observed five sequential sample draws from a hypothetical “jar” containing a certain proportion of red and blue marbles (sampling phase). The marble ratio of the unseen jar varied from trial to trial and sample draws could contain one, five or nine marbles. Afterwards, participants indicated their estimate of the proportion of blue marbles in the jar by adjusting the number of blue marbles in a 10 by 10 response grid representing the unseen jar (estimation phase; see methods for further details). Our primary measure of interest was estimation error, defined as 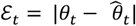, where θ is the experienced proportion of blue marbles on trial t (i.e., the mean across samples; note that this quantity differs from the true proportion of the unseen jar but constitutes the best (unbiased) estimate given the available samples) and 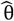 is the subject’s reported estimate of the blue marble proportion of the jar on trial t (Figure 3A). We computed the median estimation error across trials as an individual difference measure of estimation accuracy. Subjects’ median estimation error ranged from 0.01 to 0.20 (Median = 0.07, SD = 0.035). A Wilcoxon signed-ranks test revealed that estimation errors significantly differed from zero (Z = 6.215, one-sided p = 2.57 · 10^−10^). This indicates differences between experienced and estimated marble ratios on the group level. Previous research suggests that people often underestimate probabilities of frequent events and overestimate probabilities of rare events in similar decision-from-experience tasks ^47,48^. To see whether this effect contributed to estimation accuracy in our task, we examined whether subjects made more errors for jars with more extreme marble ratios. We ran a linear mixed model predicting trial-wise estimation error from the marble ratio distance from 50:50 for a given jar. To account for individual differences in intercepts, we entered subject ID as a covariate in the model. There was a significant positive relationship between trial-wise estimation-error and jar marble ratio distance from 50:50 (F(1,2702) = 154.608, p = 1.48 · 10^−34^, semi-partial η^2^ = 0.048). The more extreme the jar marble ratio, the larger subjects’ estimation error on a given trial. In the following sections, we will refer to this effect as the “extreme jar bias”. Next, we examined whether individual differences in this extreme jar bias explained individual differences in median estimation error. We computed individuals’ extreme jar bias by running regression models predicting trial-wise estimation error from marble ratio distance from 50:50 for each subject separately. Individuals’ extreme jar bias was defined as the fitted (unstandardized) slopes of these regression models (Mean = 0.198, SD = 0.230). We found that individual differences in extreme jar bias correlated strongly and positively with subjects’ median estimation error (Pearson’s r(49) = 0.63, p = 8.28 · 10^−7^; Figure 3B). Biased estimation of extreme jar proportions thus contributed to individual differences in task accuracy.

**Figure 3.**
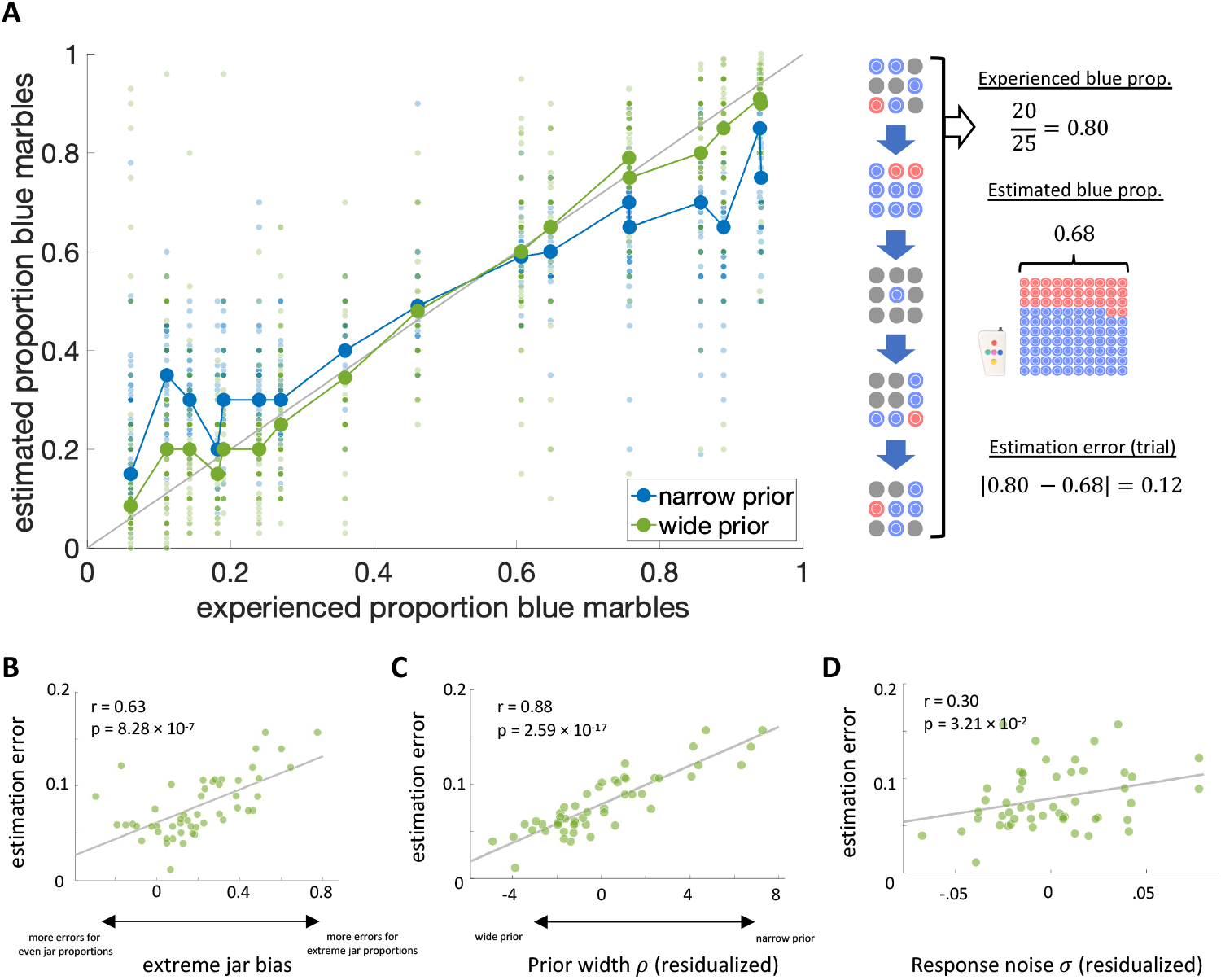
Latent sources of individual estimation errors. A: Left: Reported blue marble proportion estimates across trials and subjects against the experienced blue marble proportion across samples on a given trial. Groups are based on a median split on the fitted prior width parameter r of the Bayesian observer model (i.e., the narrow prior group had higher parameter estimates). Right: Schematic example of our trial-wise estimation error measure. B: Individual differences in median estimation error related to estimation accuracy for jars with extreme marble ratios. C-D: The fitted prior width and response noise parameters of the Bayesian observer model predicted differences in median estimation error. Plotted parameters are residualized by the other model parameter respectively.

We next fitted a set of Bayesian observer models that could account for this estimation bias in terms of inadequate uncertainty representations. The model with the overall best Bayesian Information Criterion (BIC) value was a Bayesian observer model which included a fitted prior parameter ρ, describing individual differences in the width of the belief distribution over marble ratios before observing any samples, and a response noise parameter *σ*, which captures variability of subjects’ reported marble ratio estimates around model predictions (Table 1). The second-best overall model included an exponential evidence weight *δ* and a noisy choice rule parameter σ, followed by an exponential evidence weight and fitted prior model assuming responses were sampled from the final trial posterior of the belief distribution. A Rescorla-Wagner model had the worst fit out of all models, which suggests that people indeed keep track of and leverage uncertainty estimates to update their marble ratio belief in a Bayesian manner. Parameters of the winning model were recoverable from simulations and model recovery was acceptable at the group level (see Supplementary Methods for details).

**Table 1.** Model comparison. BIC for each fitted behavioral model.

| Model | Rescorla-Wagner | Bayesian observer |  |  |  |
| --- | --- | --- | --- | --- | --- |
| Parameters | $\alpha, \sigma$ | $\rho$ | $\delta$ | $\rho, \sigma$ | $\delta, \sigma$ |
| BIC | $-3.38 \cdot 10^3$ | $-3.68 \cdot 10^3$ | $-3.82 \cdot 10^3$ | $-4.59 \cdot 10^3$ | $-4.16 \cdot 10^3$ |

We next tested whether the prior width parameter ρ of the winning Bayesian observer model captured individual differences in extreme jar bias that contributed to subjects’ estimation error while controlling for other unmodelled sources of error captured by the response noise parameter *σ*. The fitted prior parameter ρ of the winning Bayesian observer model had a median of 2.45 (SD = 4.23), indicating that subjects on average had a narrower prior than the unbiased Bayesian observer (ρ = 1). Moreover, the fitted noise parameter *σ* had a median of 0.10 (SD = 0.05), highlighting that subjects’ reported marble ratios deviated from model predictions by 10 marbles on average. We ran multiple regression analyses predicting median estimation error and extreme jar bias from the fitted prior parameter ρ and noise parameter *σ* respectively. In the first regression model, a narrower prior parameter ρ (t(48) = 13.152, p = 1.57 · 10^−17^, semi-partial η^2^ = 0.771; Figure 3C) and a higher noise parameter *σ* (t(48) = 4.501, p = 4.30 · 10^−5^, semi-partial η^2^ = 0.090; Figure 3D) uniquely predicted larger estimation errors (full model: (F(2,48) = 88.135, p = 8.54 · 10^−17^, R^2^ = 0.786). Importantly, the prior width parameter ρ accounted for most of the explained variance in estimation errors suggesting that our model captured individual differences in subjects’ response behavior well. A narrower parameter ρ reflects a stronger prior belief in a 50:50 marble ratio, which should bias estimates for more extreme jars towards this (incorrect) default belief. In line with this expectation, our second regression model revealed that subjects’ extreme jar bias was positively related to prior width ρ (t(48) = 11.780, p = 9.11 · 10^−16^, semi-partial η^2^ = 0.499) but also negatively to response noise *σ* (t(48) = -2.983, p = 4.48 · 10^−3^, semi-partial η^2^ = -0.032; full model: F(2,48) = 114.968, p = 4.956 · 10^−19^, R^2^ = 0.827). The significant (although smaller) effect of the noise parameter *σ* on extreme jar bias was driven by a small number of subjects with negative extreme jar bias slopes (i.e., subjects who made larger errors for jars with even rather than extreme marble ratios), which the Bayesian observer model could not account for (see Supplementary Results for further details). Taken together, these analyses indicate that the width of the prior belief substantially impacts individual task performance by capturing subjects’ tendency to overestimate jars with blue marble proportions close to 0 and underestimate jars with blue marble proportions close to 1 (Figure 3A).

### BOLD signal variability tracks individual differences in prior uncertainty and compresses during learning

To investigate our hypothesis that BOLD signal variability (SD_BOLD_) collapses with successive sample presentations, we ran a task PLS analysis ^45^ relating SD_BOLD_ to the five sample periods. This analysis returned a significant latent effect (permuted p = 0) showing that SD_BOLD_ reduced with successive exposure to marble samples across a distributed set of cortical brain regions spanning the parietal, prefrontal and temporal lobes (Figure 4A, see Figure S2A for full axial brain plots and Table S3 for peak voxel coordinates in robust clusters). We further investigated this trend by fitting orthogonal polynomial contrasts in a linear mixed model entering subject ID as covariate. The pattern of latent SD_BOLD_ change was best described by a linear contrast (F(1,201) = 40.444, p = 1.34 · 10^−9^, semi-partial η^2^ = 1.12 · 10^−2^) over a quadratic (F(1,201)=3.066, p = 8.15 · 10^−2^, semi-partial η^2^ = 8.41 · 10^−4^) and cubic (F(1,201)=3.416, p = 6.61 · 10^−2^, semi-partial η^2^ = 9.61 · 10^−4^) one.

**Figure 4.**
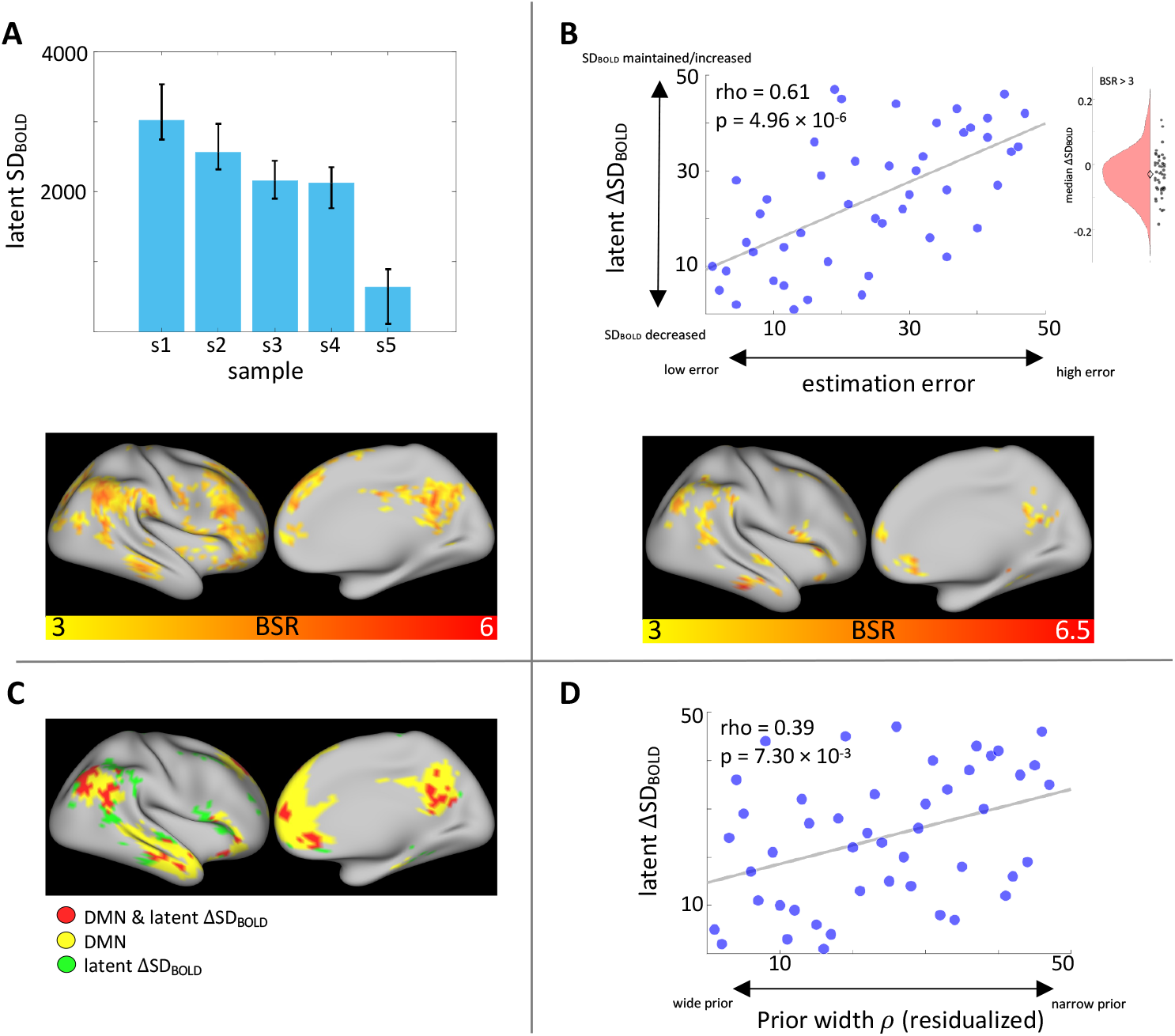
SD_BOLD_ tracks idiosyncratic uncertainty representations and predicts inference accuracy. A: Task PLS revealed average SD_BOLD_ compression over the sampling phase. Error bars represent bootstrapped 95% confidence intervals of the mean. B: Change in SD_BOLD_ over the sampling phase correlated with estimation accuracy. Greater SD_BOLD_ collapse related to smaller median estimation errors. The inset shows the distribution of median SD_BOLD_ change over positive effect voxels (BSR > 3). The diamond marker indicates the distribution median. C: Regions showing performance-related SD_BOLD_ change (see panel B) largely overlapped with the canonical Default Mode Network (DMN)^1^. D: Fitted prior width parameter estimates (residualized by noise parameter *σ*) of the Bayesian observer model positively correlated with performance-related SD_BOLD_ change. Note: All variables are rank-transformed.

Next, we asked whether the ability to collapse SD_BOLD_ over the sampling period related to individual differences in task accuracy. To this end, we quantified SD_BOLD_ change (ΔSD_BOLD_) by fitting linear regression slopes to the SD_BOLD_ estimates for the five sample presentations in each voxel. We ran behavioral PLS analysis relating these SD_BOLD_ slopes to subjects’ median estimation error. Four Cook’s d (i.e., multivariate) outliers were removed from this and all subsequent reported analyses on ΔSD_BOLD_ resulting in N = 47 (we refer to the Supplementary Information for PLS results ran on the full dataset). We found a significant latent relationship (permuted p = 3.00 · 10^−3^, Figure 4B, cf. Figure S4A for N = 51) revealing that subjects who decreased SD_BOLD_ more, especially in parietal, prefrontal (PFC) and temporal cortex, also produced smaller estimation errors (Spearman’s r = 0.612). Notably, this set of brain regions largely overlapped with the canonical Default Mode Network (DMN) that is typically observed in resting-state fMRI (BSR > 3; Figure 4C, see Figure S2B for full axial brain plots and Table S4 for peak voxel coordinates in robust clusters) ^1^.

We further investigated our hypothesis that individual differences in state uncertainty representations could explain the relationship between ΔSD_BOLD_ and estimation error. Our winning behavioral model suggested that subjects with a narrower prior belief distribution over marble ratios (i.e., large ρ parameter) showed little decrease in state uncertainty during sampling, because they started out with less uncertainty to begin with, and made more errors because they remained inflexible about their (incorrect) prior belief of a 50:50 marble ratio. In contrast, more accurate subjects (i.e., ρ close to 1) represented maximal state uncertainty at the start of a trial and reduced uncertainty more with each presented sample (Figure 2B). We expected ΔSD_BOLD_ to mirror these individual differences in state uncertainty change during the sampling phase. To test this hypothesis, we ran multiple regression analysis predicting latent ΔSD_BOLD_ (i.e., the whole-brain pattern of ΔSD_BOLD_ which relates to median estimation error; see Figure 3b) from the prior width ρ parameter of the winning behavioral model. We also entered the response noise parameter *σ* as a control variable, which captures all unmodeled sources of estimation error and thus serves as a measure of model misfit (see Methods). This model explained a significant amount of variance (F(2,44) = 4.489, p = 1.68 · 10^−2^, R^2^ = 0.169). There was a significant main effect only for prior width ρ (t(44) = 2.703, p = 9.72 · 10^−3^, semi-partial η^2^ = 0.138) but not for noise parameter *σ* (t(44) = 0.425, p = 0.673). Therefore, the wider a participant’s prior belief (i.e., smaller ρ), the more they collapsed SD_BOLD_ during the sampling phase (Figure 4D). To ensure that that these effects were not due to the specific parameterization of the winning behavioral model, we replicated this analysis also for the parameters of the (second-best) evidence weight model (see Supplementary Results).

Overall, these results indicate that the degree to which subjects compress SD_BOLD_ with learning predicts individual differences in inference accuracy, which can be explained in terms of idiosyncratic uncertainty representations.

#### SD_BOLD_ compression effects were unrelated to interindividual SD_BOLD_ levels

It is possible that our within-person effects of ΔSD_BOLD_ are not independent of between-subject differences in SD_BOLD_ levels, which have also been linked to task performance in previous studies ^13^. To account for this, we computed a latent SD_BOLD_ control variable by matrix multiplying brain saliences from the behavioral PLS analysis relating ΔSD_BOLD_ to estimation error (which reflect the spatial expression of the observed relationship) with subjects’ SD_BOLD_ level at the first sample presentation period. We found no significant correlation between latent ΔSD_BOLD_ and latent SD_BOLD_ at first sample exposure (Spearman’s r(45) = -0.073, p = 6.28 · 10^−1^). Next, estimation error was significantly related to ΔSD_BOLD_ (t(44) = 5.430, p = 2 · 10^−6^, semi-partial η^2^ = 0.625) when controlling for the effect of latent SD_BOLD_ at first sample exposure (t(44) = 1.806, p = 7.78 · 10^−2^, semi-partial η^2^ = 0.208; full model: F(2,44) = 15.747, p = 7 · 10^−2^, R^2^ = 0.417). Likewise, prior width ρ (residualized by noise parameter *σ*) of the winning Bayesian-observer model was significantly related to ΔSD_BOLD_ (t(44) = 3.151, p = 2.92 · 10^−3^, semi-partial η^2^ = 0.409) when controlling for the effect of latent SD_BOLD_ at first sample exposure (t(44) = 2.526, p = 1.52 · 10^−2^, semi-partial η^2^ = 0.328; full model: F(2,44) = 7.618, p = 1.44 · 10^−3^, R^2^ = 0.257). In both models, the inclusion of the control variable also did not diminish the effect size of ΔSD_BOLD_ on these behavioral measures. Thus, the effect of within-person SD_BOLD_ compression on task performance was independent of between-subject differences in initial SD_BOLD_ levels.

#### SD_BOLD_ compression effects were unrelated to change in BOLD response magnitude

Finally, we investigated whether our SD_BOLD_ findings were related to change in average BOLD response magnitude over the sampling phase. To this end, we obtained voxel-wise GLM beta estimates for each sample period (MEAN_BOLD_) and entered them into a task PLS. This analysis returned a significant latent effect (permuted p = 0) showing that MEAN_BOLD_ increased from the first to last sample presentation in large parts of the cortex (including occipital, parietal, and pre-/frontal regions) and subcortex (mainly thalamus and striatum; Figure S3, Table S5). The pattern of latent MEAN_BOLD_ change was best described by a cubic contrast (F(1,201) = 17.984, p = 3.40 · 10^−5^, semi-partial η^2^ = 4.91 · 10^−2^). We then ran a behavioral PLS analysis to examine whether interindividual differences in MEAN_BOLD_ change related to task performance (i.e., subjects’ median estimation errors). In line with the change trajectory suggested by our task PLS results, we first quantified MEAN_BOLD_ change by fitting a cubic regression coefficient to the MEAN_BOLD_ estimates for the five sample presentations in each voxel and subject. Unlike for SD_BOLD_ (see Fig 4B), there was no significant latent relationship (permuted p = 0.173 for N = 48 with Cook’s d outliers removed; permuted p = 0.755 for N = 51). To ensure that this lack of association to subjects’ estimation error was not dependent on fitting a cubic change trajectory, we also ran two additional PLS analyses quantifying MEAN_BOLD_ change by a fitted linear and quadratic regression coefficient respectively. Neither PLS analysis detected a relationship between MEAN_BOLD_ change and subjects’ estimation error (linear: permuted p = 0.321 for N = 50 with Cook’s d outliers removed; permuted p = 0.275 for N = 51); quadratic: permuted p = 0.258 for N = 47 with Cook’s d outliers removed; permuted p = 0.474 for N = 51).

Finally, we directly assessed whether the observed association between ΔSD_BOLD_ and task performance would still hold when controlling for MEAN_BOLD_ change. To this end, we obtained a latent variable of MEAN_BOLD_ change by (1) taking the brain saliences (i.e., spatial weights) from the behavioral PLS analysis on ΔSD_BOLD_ (see Fig. 4B) and then (2) computing the dot product of those weights with MEAN_BOLD_ change data for each subject. This latent variable allowed for a spatially-matched comparison of SD_BOLD_ and MEAN_BOLD_ change in the same brain regions that characterized the ΔSD_BOLD_ effect shown in Fig 4B. We carried out these steps three times for the fitted linear, quadratic and cubic regression coefficient of MEAN_BOLD_ change respectively, thus taking into account different possible shapes of MEAN_BOLD_ change trajectories. Three separate regression models consistently revealed a significant main effect for ΔSD_BOLD_ on estimation error, even when controlling for (1) linear MEAN_BOLD_ change (SD_BOLD_: t(44) = 5.158, p = 6.00 · 10^−6^, semi-partial η^2^ = 0.377; MEAN_BOLD_: t(44) = -0.444, p = 6.59 · 10^−1^, semi-partial η^2^ = 0.003), (2) quadratic MEAN_BOLD_ change (SD_BOLD_: t(44) = 5.157, p = 6.00 · 10^−6^, semi-partial η^2^ = 0.376; MEAN_BOLD_: t(44) = 0.527, p = 6.01 · 10^−1^, semi-partial η^2^ = 0.004), or (3) cubic MEAN_BOLD_ change (SD_BOLD_: t(44) = 5.383, p = 3.00 · 10^−6^, semi-partial η^2^ = 0.396; MEAN_BOLD_: t(44) = -1.371, p = 1.77 · 10^−1^, semi-partial η^2^ = 0.026). Together, these analyses confirm that only modulation of SD_BOLD_ but not MEAN_BOLD_ was predictive of subjects’ inference accuracy.

### Standard GLM neural uncertainty correlates are distinct from BOLD variability effects

Notably, change in MEAN_BOLD_ over the sampling period does not account for more nuanced BOLD response modulation by uncertainty resulting from the varying sample sizes in our task design (see Figure S1B). In other words, an unbiased Bayesian observer model would predict more uncertainty reduction after observing a larger compared to a smaller sample in addition to the overall reduction in uncertainty with the total accumulated evidence (i.e., trial time). Our SD_BOLD_ metric required discrete uncertainty conditions (with sufficient trial counts) and we thus only considered the main effect of trial time on uncertainty. However, in a standard GLM approach we could model the parametric modulation of the BOLD signal by the posterior variance of the belief distribution (i.e., uncertainty) for the unbiased Bayesian observer model, which accounts for both trial time and sample size effects on uncertainty. Again, we used a multivariate PLS approach to relate individuals’ voxel-wise GLM beta estimates (for a parametric uncertainty regressor) to median estimation error. This revealed a significant latent association (permuted p = 3.40 · 10^−2^, Spearman’s r = 0.675, Figure 5A, cf. Figure S4B for N = 51). Robust positive clusters were mainly located in the lateral parietal cortex, PFC (dorsomedial, ventrolateral, and rostrolateral parts), and in the insula. Robust negative clusters were present in ventromedial PFC (vmPFC) and in the left hippocampus (see Figure S2C for full axial brain plots and Table S6 for peak voxel coordinates in robust clusters). This latent relationship was related to the prior width parameter ρ of the winning Bayesian observer model, similar to the effect of ΔSD_BOLD_ (Figure 5B). For further details on these results, please refer to the Supplementary Information.

**Figure 5.**
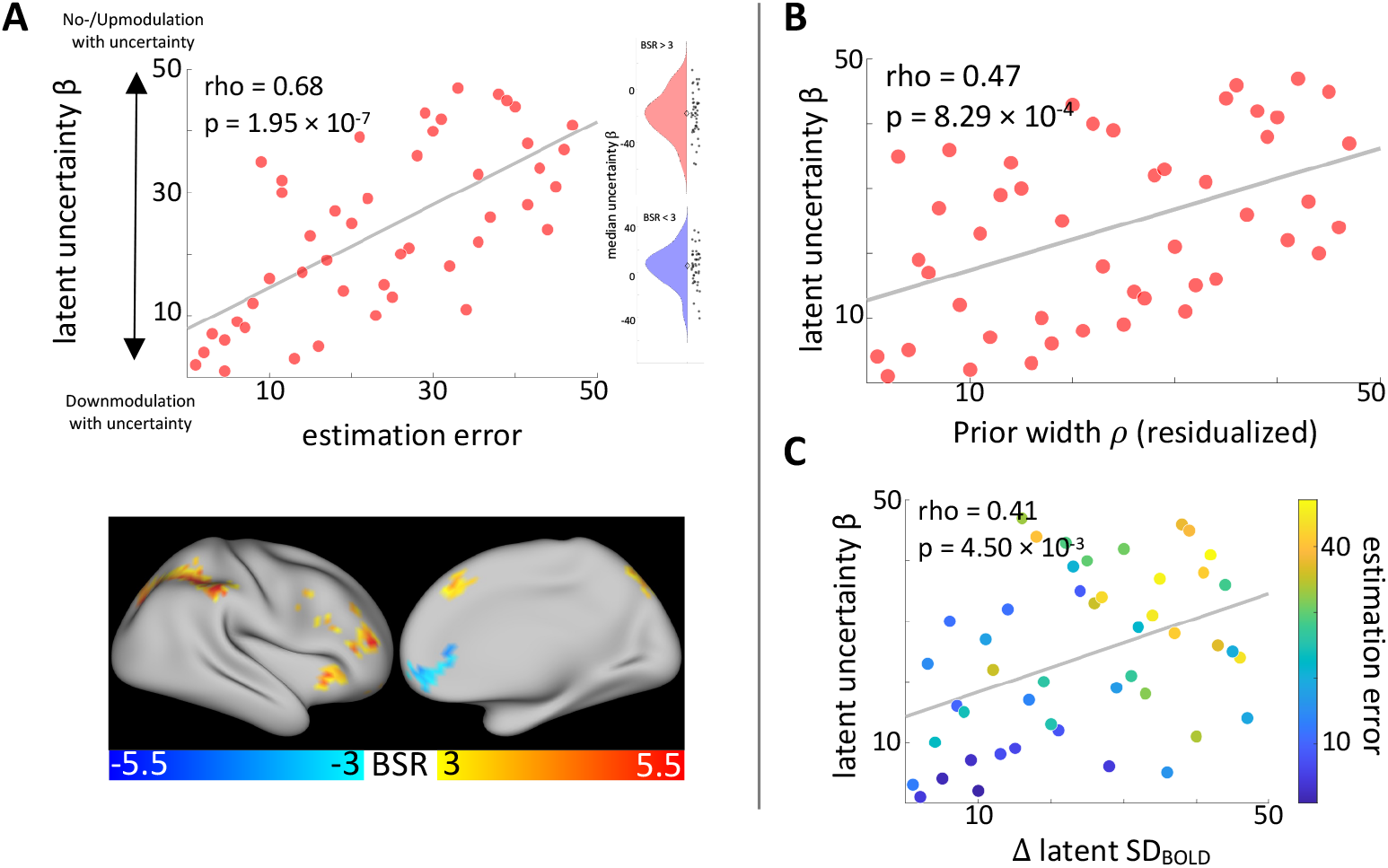
Standard GLM neural uncertainty correlates are distinct from SD_BOLD_ effects. A: BOLD signal modulation by uncertainty (GLM b estimates) correlated with average estimation accuracy. More downmodulation with increasing state uncertainty across the whole brain related to smaller estimation errors. The inset shows the distribution of b estimates over positive (BSR > 3, top) and negative (BSR < 3, bottom) effect voxels. The diamond marker indicates the distribution median. B: Fitted prior width parameter values (residualized by noise parameter *σ*) of the Bayesian observer model positively correlated with performance-related BOLD signal modulation by uncertainty (GLM b estimates). C: Changes in SD_BOLD_ over the sampling phase and BOLD signal modulation by uncertainty (GLM b estimates) uniquely predicted task performance. Good performers showed both more SD_BOLD_ compression and more downmodulation of the BOLD signal when state uncertainty was high. Note: All PLS results are presented for rank-transformed variables.

Qualitatively, these effect regions seem to show little overlap with those showing an effect for ΔSD_BOLD_. To test the spatial specificity of ΔSD_BOLD_ effects, we computed a new latent variable by extracting the brain saliences from the behavioral PLS analysis relating latent ΔSD_BOLD_ to estimation error and matrix-multiplied them with the uncertainty beta estimates in each voxel from our standard parametric GLM analysis. We then ran multiple regression analysis predicting median estimation error from latent ΔSD_BOLD_ and this spatially-matched latent parametric uncertainty modulation variable. The regression model was significant (F(2,44) = 13.901, p = 2.10 · 10^−5^) with an R^2^ of 0.387. The main effect of latent ΔSD_BOLD_ was significant (t(44) = 5.265, p = 4 · 10^−6^, semi-partial η^2^ = 0.386) but not for the new latent control variable (t(44) = 0.975, p = 0.335). This indicates that ΔSD_BOLD_ effects and the parametric uncertainty modulation of the BOLD signal on performance are spatially distinct in the brain.

Furthermore, we found a significant positive correlation between the latent ΔSD_BOLD_ (Spearman’s r(45) = 0.407, p = 4.55 · 10^−3^) and latent uncertainty modulation effects on estimation accuracy indicating that more accurate performers show both types of uncertainty representation in the brain. We ran multiple regression analysis predicting estimation errors from both latent ΔSD_BOLD_ and latent uncertainty modulation to investigate whether they explained unique variance in estimation errors. The full model was significant (F(2,44) = 31.883, p = 2.76 · 10^−9^, R^2^ = 0.592; Figure 5C). We found a significant main effect of both latent ΔSD_BOLD_ (t(44) = 4.844, p = 1.60 · 10^−5^, semi-partial η^2^ = 0.218) and latent uncertainty modulation (t(44) = 3.829, p = 4.04 · 10^−4^, semi-partial η^2^ = 0.136). This suggests that both types of neural uncertainty correlates uniquely relate to task accuracy.

Overall, these analyses reveal that SD_BOLD_ effects are spatially distinct and uniquely relate to task accuracy compared to neural correlates revealed by a more standard GLM approach.

## DISCUSSION

In this study we show that reductions in state uncertainty over the course of learning co-occur with a compression in BOLD signal variability. Moreover, individuals who decreased state uncertainty to a greater extent showed more pronounced SD_BOLD_ compression and made smaller estimation errors on average. Behavioral modeling suggested that better performers reduced state uncertainty to a greater extent because they began with a wider (more uncertain and flexible) prior belief distribution before observing any samples, resulting in more unbiased marble ratio estimates.

Our findings establish SD_BOLD_ as a novel within-person neural correlate of uncertainty, which reduces over the course of Bayesian inference/learning, and relates to inference accuracy. The current findings thus add to a growing literature demonstrating that within-person SD_BOLD_ modulation in the face of varying task demands facilitates adaptive behavior across different cognitive domains (see ^13^). Our study is the first to directly show that SD_BOLD_ tracks reductions in uncertainty. The bulk of past empirical work arguing for such a link showed modulation of brain signal variability across disparate task conditions that varied along several dimensions beyond uncertainty (e.g., cognitive demand, bottom-up sensory input or processing requirements ^17,18,49,50^). In contrast, here we employed a learning paradigm, which enabled us to operationalize uncertainty in a precise manner; specifically, one’s uncertainty regarding the latent state of an environmental variable, a property that can be reduced through learning from observations. We used this paradigm in conjunction with a novel event-related method to quantify SD_BOLD_, which allowed us to track the evolution of SD_BOLD_ with changes in uncertainty on a short time scale (several seconds). This method was inspired by the machine-learning fMRI literature ^44^ and is a stark departure from previous fMRI work, which usually computed SD_BOLD_ over entire task blocks ^13,14^. Moreover, our control analyses revealed that, unlike SD_BOLD_, modulation of mean BOLD response magnitude with each newly presented sample was not predictive of subject’s inference accuracy and did not diminish our SD_BOLD_ effects when statistically controlled. This result was expected given that previous studies have reported only modest associations between SD_BOLD_ and mean BOLD response magnitude ^15,41,51,52^.

Due to the limited temporal resolution of fMRI (which was one time point for each sample period on a given trial in our study design), we defined SD_BOLD_ as the trial-to-trial variability in the BOLD signal from each within-trial sampling period, which we have argued serves as a direct proxy for within-trial neural dynamics (see Fig. 1). It is important to note that in our task design, bottom-up visual input and difficulty was matched across trials. Therefore, we believe the key source of variability contributing to across-trial BOLD variability is within-trial neural variability related to stimulus processing (i.e., learning/uncertainty). Our findings thus align well with recent evidence showing that (within-trial) EEG signal entropy decreases with decreasing uncertainty on a perceptual decision-making task, which also matched bottom-up visual inputs between task conditions ^12^.

Taken together, our results thus support the longstanding idea that more brain signal variability enables more cognitive flexibility ^13,14^, which we show is required under high uncertainty early on in the learning process to efficiently update one’s internal belief about the environmental state.

### Individual differences in uncertainty representations track inference accuracy

We found that low performing individuals made most estimation errors for extreme marble ratios, such that jars with blue marble proportions close to one were underestimated and those with proportions close to zero were overestimated. This effect has been reported previously for tasks where participants are asked to judge outcome probabilities from experience, sometimes referred to as under-extremity ^47,48,53^. Notably, some studies found that this effect is abolished when participants were asked to give verbal judgements instead ^54^. This may suggest that individual differences in extreme jar bias result from people’s (in)ability to accurately enter their marble ratio estimates into the grid in the estimation phase of our task. However, we found that brain activity during sampling (*prior* to the estimation phase) predicted individual differences in peoples’ (average) estimation error. It is thus likely that the observed bias reflects people’s internal representations rather than being a mere consequence of our chosen response modality.

Our winning Bayesian observer model suggested that such estimation errors could best be explained by individual differences in the representation of the initial prior belief over blue marble proportions rather than in belief updating; low performers held a narrower prior belief distribution over the default belief of a 50:50 marble ratio. It is possible that this prior was induced by our task design given the default position of the response grid. The fitted prior could thus reflect individual differences in anchoring to this task feature ^55^. Alternatively, this bias could reflect bona fide trait-like individual differences in expectations about extreme outcomes. In line with this explanation, Glaze, et al. ^56^ found that performance differences on a change-point inference task were best explained by participants’ prior width over potential change rates, which cannot simply be explained by anchoring. Note that the distinction between prior beliefs and belief updating is also pertinent to other approximately-Bayesian and non-Bayesian cognitive models that have been used to model the inference process in similar tasks, which we did not consider here ^48,57^. These models account for cognitive limitations in human information processing and may provide an even better fit to behavioral data. While not the main aim of the current study, future modelling and experimental work is required to pin down the source of individual differences in under-extremity observed in the current study.

We showed that less SD_BOLD_ compression (independent of initial SD_BOLD_ levels) in poorer performers could best be explained by more limited uncertainty reduction, based on parameter estimates for the winning behavioral model. Still, our results rely on the assumption that participants represent and utilize uncertainty akin to an (approximate) Bayesian observer. Future work should reduce reliance on model assumptions and obtain subjective uncertainty estimates. For example, one could regularly ask people to report their state belief together with their belief confidence throughout the learning process to directly track updates in their subjective belief distribution.

### Performance-related SD_BOLD_ collapse in the Default Mode Network

We observed performance-relevant collapse in SD_BOLD_ in a network of brain regions that largely overlapped with regions commonly ascribed to the default mode network (DMN) in resting-state fMRI connectivity analyses ^1,58-61^, including parietal (precuneus, inferior parietal lobe including the angular gyrus), prefrontal (superior frontal gyrus, frontopolar cortex, orbitofrontal cortex, paracingulate gyrus) and temporal (middle temporal gyrus) cortices. Previous work has consistently reported deactivation of this brain network during externally-cued, demanding cognitive tasks compared to rest ^58,59,61-64^. In contrast, internally-oriented tasks that call for self-referential processing and memory recollection demonstrate increased activity in key regions of the DMN ^61,65-67^. With respect to brain signal variability, one study by Grady and Garrett ^49^ found *increased* SD_BOLD_ in DMN regions in externally-as compared to internally-guided tasks. Given previous reports that higher local SD_BOLD_ is associated with lower whole-brain functional network dimensionality ^68^, this finding may reflect more crosstalk between the DMN and other brain networks during externally-guided tasks. Therefore, SD_BOLD_ compression during learning may reflect the gradual build-up of an internal world model; larger SD_BOLD_ under high state uncertainty early on may afford a more flexible incorporation of incoming information to update one’s internal state belief. As state uncertainty reduces and SD_BOLD_ compresses, the brain arrives at a stable internal belief representation, which informs one’s jar marble ratio estimate in the subsequent task phase.

Previous research has shown that internally-guided tasks evoke a coupling between the DMN and the frontoparietal-control network to support task performance ^69^. In our task, we would expect such a coupling to emerge towards the end of the sampling period reflecting the utilization of one’s internal state belief to prepare the upcoming response. Indeed, we observe a concomitant increase in BOLD activity in the fronto-parietal network and compression of SD_BOLD_ in the DMN in high performing subjects. How functional connectivity between different brain networks changes over the course of learning goes beyond the scope of the current study, but constitutes an interesting target for future research.

### Standard analytic approaches reveal different neural uncertainty correlates compared to SD_BOLD_

Our standard GLM analysis revealed that BOLD modulation by uncertainty, as derived from an unbiased Bayesian observer model, also related to estimation accuracy in our task. On one hand, higher BOLD (in lateral parietal cortex, dorsomedial PFC, ventro- and rostrolateral PFC, and anterior insula) with lower uncertainty predicted higher estimation accuracy. In other words, high performers showed an increase in BOLD activity in these regions as they observed more samples and became more certain about the jar marble ratio on a given trial. These areas correspond to fronto-parietal control and dorsal attention brain networks, which support goal-directed behavior in externally-driven tasks ^1,70-73^. Conversely, *positive* coupling between the BOLD signal and state uncertainty in mainly in the vmPFC (and a cluster in the left hippocampus) also predicted better performance in our task, which has previously been linked to confidence representations ^74^ (see Supplementary Information for detailed discussion of these findings).

Interestingly, we found that uncertainty representations in SD_BOLD_ (in DMN regions) and parametric BOLD signal modulation (in fronto-parietal control/attention regions) were spatially different and *uniquely* predicted estimation accuracy. Why would the brain track uncertainty in these different neural signals and across different networks? One possibility could be that SD_BOLD_ tracks uncertainty about the latent environmental state with or without immediate relevance for decision-making, while parametric BOLD modulation by uncertainty may be more relevant in the context of goal-directed decision formation and action selection. In support of this view, previous research has shown modulation of brain signal variability by stimulus features even when no action was required. For example, in the human neuroimaging literature, changes in neural variability have been observed in response to varying complexity of visual stimuli during passive viewing, which were linked to offline cognitive performance ^46,50^. In a similar vein, it has recently been argued that early perceptual uncertainty, which has been the main focus of sampling accounts of neural variability, is tracked irrespective of task demands but can be flexibly utilized in higher-order decision-making ^6-8,75^. Thus, changes in brain signal variability may track environmental variables without the need for immediate action. In contrast, parametric BOLD modulation by uncertainty has been reported for learning and decision-making tasks that require participants to make choices on every trial ^2,23-37^. As such, these neural uncertainty correlates may be more tightly linked to decision formation and may in some cases even relate to metacognitive awareness of this decision-relevant variable ^74,76,77^. Although speculative, this framework makes testable predictions for situations in which we would expect to see a dissociation of these two neural signals, which should be addressed in future work.

### Limitations and next steps

One potential limitation of our task design was that our stimulus sample size manipulation introduced trial-to-trial variance in how much uncertainty could be reduced with each presented sample (Figure S1B). Because our SD_BOLD_ measure was computed for each sample period across trials, uncertainty levels were by definition not equivalent within each condition bin (although uncertainty always decreased across sample periods). However, our results showed that SD_BOLD_ effects could not be explained by parametric BOLD signal modulation by trial-to-trial differences in uncertainty trajectories in the same brain regions, which could have accounted for this variance. Ideally, future studies should keep sample size consistent between trials so that SD_BOLD_ can be computed over comparable uncertainty levels.

Another potential limitation is that uncertainty was collinear with within-trial time in our task; the more samples were presented, the more uncertainty could be reduced in general. However, this is an inevitable consequence of decision-making and learning in stable environments, in which one can simply accumulate evidence over time. Although within-trial time could have introduced arousal- or attention-related effects (which have previously been linked to human brain signal variability ^13^), we observed performance-related interindividual variability in SD_BOLD_ compression over the sampling period. Such an effect cannot, by definition, be accounted for by the fixed factor of trial time. Nevertheless, future work could investigate the coupling between brain signal variability and uncertainty in non-stationary learning environments (see e.g. ^3^), within which uncertainty can increase *or* decrease over time.

## Conclusion

We provide first evidence that moment-to-moment brain signal variability compresses with increasing belief precision during learning. Whether brain signal compression is directly proportional to the informativeness of the available evidence is an important prediction that should be investigated in future work.

## Acknowledgements and Funding

During the work on his dissertation, A.S. was a pre-doctoral fellow of the International Max Planck Research School on the Life Course (LIFE, www.imprs-life.mpg.de; participating institutions: Max Planck Institute for Human Development, Freie Universität Berlin, Humboldt-Universität zu Berlin, University of Michigan, University of Virginia, University of Zurich) and the International Max Planck Research School on Computational Methods in Psychiatry and Ageing Research (COMP2PSYCH, https://www.mps-ucl-centre.mpg.de/comp2psych). D.D.G. received an Emmy Noether Programme grant from the German Research Foundation. D.D.G. is affiliated with the Max Planck UCL Centre for Computational Psychiatry and Aging Research.

