## Supplementary material for "Neural variability compresses with increasing belief precision during Bayesian inference": Supp material

### SUPPLEMENTARY INFORMATION

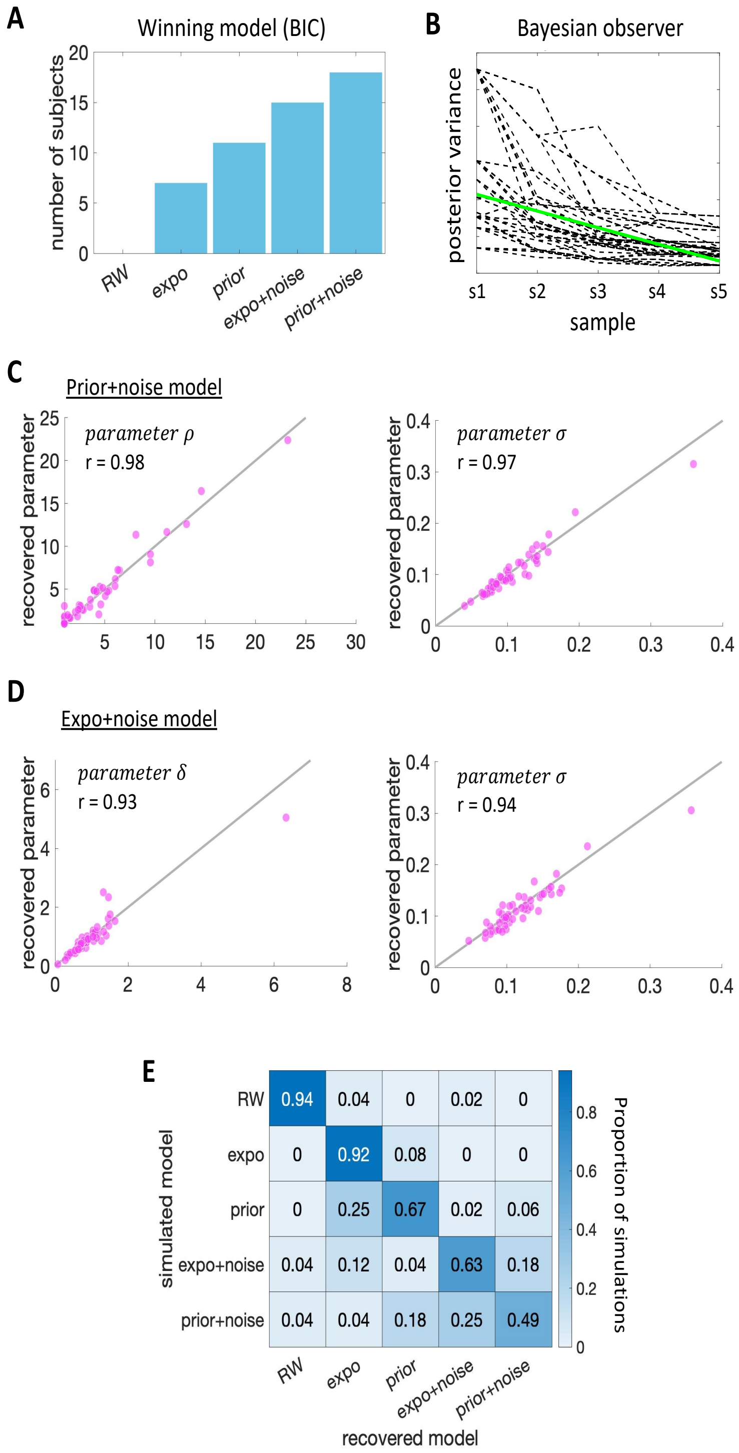


*Figure S1.* Model comparison and simulations. A: Counts of models with the lowest BIC across subjects. B: Trial-wise trajectories of the posterior variances of the marble ratio belief distribution for the unbiased Bayesian observer model. C: Parameter recovery for the Bayesian observer model with free prior width $\rho$ and noisy choice $\sigma$. D: Parameter recovery for the Bayesian observer model with free evidence weight $\delta$ and noisy choice $\sigma$. E: Proportion of models recovered from simulated subjects for each model. *Note.* RW = Rescorla-Wagner model, expo = Bayesian observer with exponential evidence weight $\delta$ and sampling choice rule, prior = Bayesian observer model with fitted prior width $\rho$ and sampling choice rule, expo+noise = same as expo model but with noisy choice $\sigma$, prior+noise = same as prior but with noisy choice $\sigma$

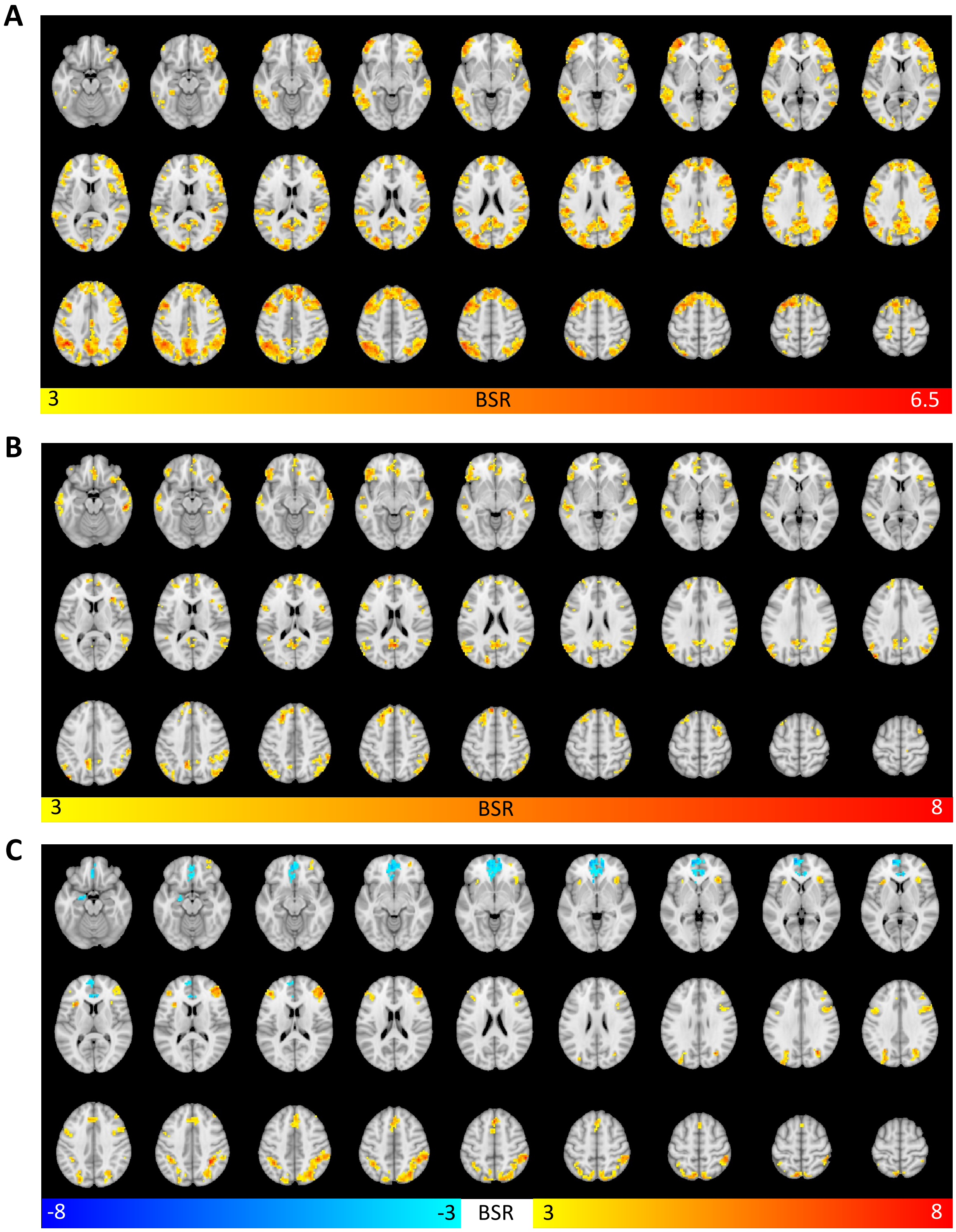


*Figure S2.* Axial brain plots for the PLS results. A: Task PLS model relating $SD_{BOLD}$ to sample presentations. B: Behavioral PLS model relating $\Delta SD_{BOLD}$ to median estimation error (N = 47). C: Behavioral PLS model relating BOLD signal uncertainty modulation to median estimation error (N = 47). Bootstrap ratio = BSR.

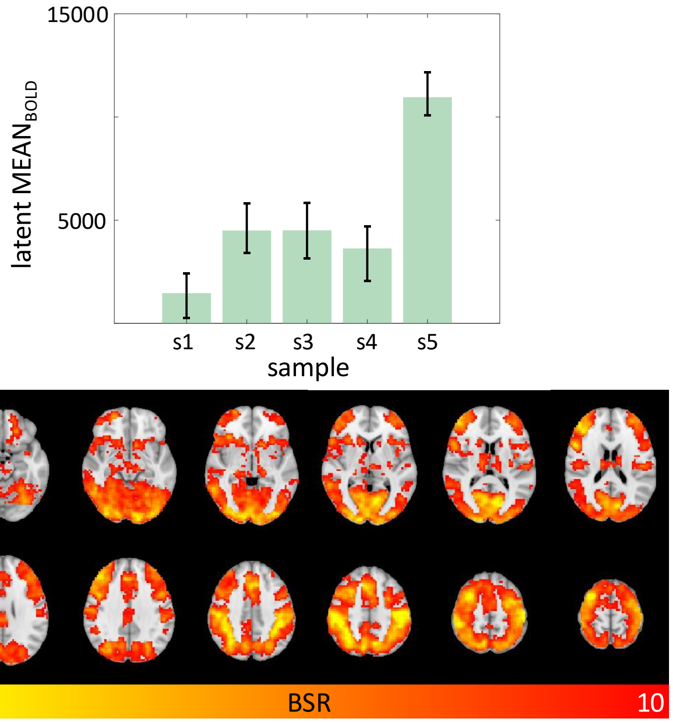


*Figure S3.* Task PLS revealed average $MEAN_{BOLD}$ increased over the sampling phase. Error bars represent bootstrapped 95% confidence intervals of the mean. Note: The results are presented for rank scored variables.

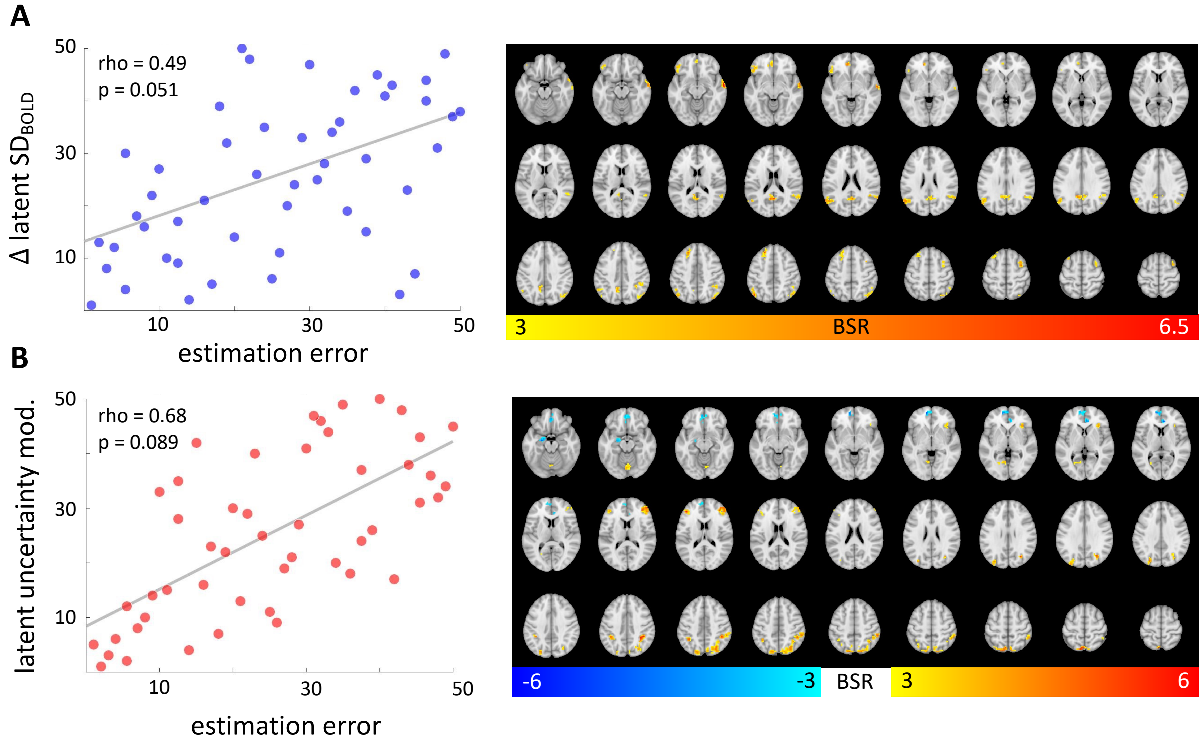


*Figure S4.* Behavioral PLS results for the full dataset (N = 51). A: Behavioral PLS model relating $\Delta SD_{BOLD}$ to average estimation error. B: Behavioral PLS model relating BOLD signal uncertainty modulation to median estimation error. Note: The results are presented for rank scored variables. P-values are based on nonparametric permutation tests.

***Supplementary Table S1.*** Overview of the 18 marble jars. Each participant completed each jar four times (twice in the low reward and twice in the high reward condition).

| Jar | Jar type | p(blue) | Payoff gamble | EV gamble | EV certain | EV difference |
|---|---|---|---|---|---|---|
| **1** | risk seeking | 0.2 | 55 | 11 | 10 | 1 |
| **2** | risk seeking | 0.1 | 130 | 13 | 10 | 3 |
| **3** | risk seeking | 0.4 | 28 | 11.2 | 10 | 1.2 |
| **4** | risk seeking | 0.6 | 22 | 13.2 | 10 | 3.2 |
| **5** | risk seeking | 0.9 | 12 | 10.8 | 10 | 0.8 |
| **6** | risk seeking | 0.8 | 16 | 12.8 | 10 | 2.8 |
| **7** | risk neutral | 0.1 | 100 | 10 | 10 | 0 |
| **8** | risk neutral | 0.2 | 50 | 10 | 10 | 0 |
| **9** | risk neutral | 0.25 | 40 | 10 | 10 | 0 |
| **10** | risk neutral | 0.77 | 13 | 10 | 10 | 0 |
| **11** | risk neutral | 0.83 | 12 | 10 | 10 | 0 |
| **12** | risk neutral | 0.91 | 11 | 10 | 10 | 0 |
| **13** | risk averse | 0.1 | 92 | 9.2 | 10 | -0.8 |
| **14** | risk averse | 0.2 | 35 | 7 | 10 | -3 |
| **15** | risk averse | 0.6 | 15 | 9 | 10 | -1 |
| **16** | risk averse | 0.4 | 17 | 6.8 | 10 | -3.2 |
| **17** | risk averse | 0.8 | 11 | 8.8 | 10 | -1.2 |
| **18** | risk averse | 0.9 | 8 | 7.2 | 10 | -2.8 |

*Note.* EV = Expected value, p(blue) = blue marble probability

***Supplementary Table S2.*** Model recovery. BIC of each model for each simulated dataset.

| **Recovered model:** | **RW** | **expo** | **prior** | **expo+noise** | **prior+noise** |
|---|---|---|---|---|---|
| **RW** | -3508.163 | -1984.135 | 1560.267 | -2694.659 | -2633.512 |
| **expo** | -2828.659 | -4926.453 | -3708.676 | -4223.386 | -4018.178 |
| **prior** | -3706.85 | -5704.678 | -6375.963 | -5297.432 | -5702.832 |
| **expo+noise** | -2888.764 | -3287.136 | -2035.45 | -4396.865 | -4037.069 |
| **prior+noise** | -3207.927 | -3669.239 | -3177.083 | -4293.817 | -4770.482 |

**Supplementary Table S3**: *Peak voxel coordinates in robust clusters (BSR > 3) for the significant latent variable of the task PLS analysis relating $SD_{BOLD}$ to sample presentation.*

| | MNI | | | | | |
|---|---|---|---|---|---|---|
| **X** | **Y** | **Z** | **BSR** | **p-value** | **Cluster size (voxels)** | **Anatomical region** |
| -39 | -54 | 36 | 6.4734 | 0.0000 | 4086 | L Inferior Parietal Lobule |
| -33 | 51 | 0 | 6.3604 | 0.0000 | 488 | L Middle Frontal Gyrus |
| -45<br>(-45) | 12<br>(9) | 54<br>(54) | 6.2607 | 0.0000 | 3370 | L Middle Frontal Gyrus* |
| -9 | -96 | 12 | 5.9355 | 0.0000 | 384 | L Superior Occipital Gyrus |
| 60 | -24 | -3 | 5.2715 | 0.0000 | 189 | R Middle Temporal Gyrus |
| 3 | -6 | 66 | 4.5180 | 0.0000 | 36 | R Posterior-Medial Frontal |
| -30 | -39 | -12 | 4.5173 | 0.0000 | 25 | L Fusiform Gyrus |
| 15 | 48 | 0 | 4.3964 | 0.0000 | 58 | R Superior Medial Gyrus |
| 45 | -6 | -3 | 4.2319 | 0.0000 | 27 | R Insula Lobe |
| -24 | -39 | 66 | 4.1320 | 0.0000 | 46 | L Postcentral Gyrus |
| 18<br>(22) | -24<br>(-24) | 63<br>(63) | 4.0587 | 0.0000 | 52 | R Precentral Gyrus* |

*Note.* Abbreviations: MNI = Coordinates in Montreal Neurological Institute space, BSR = Bootstrap ratio. Anatomic labels were derived from the cytoarchitectonic atlas provided in the anatomy toolbox for SPM 8 (Eickhoff et al., 2005). We only report clusters of 25 voxels or more. *Peak voxel not assigned anatomical label in anatomy toolbox. Anatomical label corresponds to bracketed coordinates.

**Supplementary Table S4**: *Peak voxel coordinates in robust clusters (BSR > 3) for the significant latent variable of the behavioral PLS analysis relating ΔSD$_{BOLD}$ to estimation error (N = 47).*

| | MNI | | | | | |
|---|---|---|---|---|---|---|
| **X** | **Y** | **Z** | **BSR** | **p-value** | **Cluster size (voxels)** | **Anatomical region** |
| 57 | 3 | -24 | 7.6886 | 0.0000 | 180 | R Middle Temporal Gyrus |
| 3 | -57 | 18 | 7.5207 | 0.0000 | 257 | R Precuneus |
| -45 | -78 | 33 | 7.4781 | 0.0000 | 275 | Area PGp (IPL) |
| -21 | 33 | 48 | 7.3272 | 0.0000 | 187 | L Superior Frontal Gyrus |
| 33 | 27 | 9 | 6.6384 | 0.0000 | 26 | R Insula Lobe |
| 48 | -69 | 33 | 5.9344 | 0.0000 | 180 | R Angular Gyrus |
| -9 | 66 | 18 | 5.8743 | 0.0000 | 178 | L Superior Medial Gyrus |
| -39 | 42 | -6 | 5.8357 | 0.0000 | 169 | L IFG (p. Orbitalis) |
| 60 | -45 | 48 | 5.7765 | 0.0000 | 327 | R Inferior Parietal Lobule |
| -15 | -87 | 21 | 5.7644 | 0.0000 | 32 | L Superior Occipital Gyrus |
| -54 | -27 | -3 | 5.7185 | 0.0000 | 51 | L Middle Temporal Gyrus |
| 36 | 3 | 57 | 5.5382 | 0.0000 | 84 | R Middle Frontal Gyrus |
| 51 | 6 | 15 | 5.5042 | 0.0000 | 107 | R Rolandic Operculum |
| 36 | 18 | -18 | 5.4959 | 0.0000 | 42 | R Insula Lobe |
| 12 | -24 | 69 | 5.2755 | 0.0000 | 39 | R Paracentral Lobule |
| 24 | -36 | -6 | 5.1858 | 0.0000 | 25 | R ParaHippocampal Gyrus |
| -9 | 57 | 39 | 5.1066 | 0.0000 | 53 | L Superior Frontal Gyrus |
| -45 | -45 | 15 | 4.9372 | 0.0000 | 51 | L Superior Temporal Gyrus |
| -51 | 12 | 18 | 4.9333 | 0.0000 | 43 | L IFG (p. Opercularis) |
| -66 (-63) | -27 (-31) | -24 (-20) | 4.8241 | 0.0000 | 133 | L Inferior Temporal Gyrus* |
| 15 | 36 | 51 | 4.8089 | 0.0000 | 45 | R Superior Frontal Gyrus |
| -33 | 60 | 18 | 4.6589 | 0.0000 | 50 | L Middle Frontal Gyrus |
| 9 | 60 | 15 | 4.5953 | 0.0000 | 54 | R Superior Medial Gyrus |
| 42 | 57 | 15 | 4.1974 | 0.0000 | 70 | R Middle Frontal Gyrus |

*Note.* Abbreviations: MNI = Coordinates in Montreal Neurological Institute space, BSR = Bootstrap ratio. Anatomic labels were derived from the cytoarchitectonic atlas provided in the anatomy toolbox for SPM 8 (Eickhoff et al., 2005). We only report clusters of 25 voxels or more. *Peak voxel not assigned anatomical label in anatomy toolbox. Anatomical label corresponds to bracketed coordinates.

**Supplementary Table S5**: *Peak voxel coordinates in robust clusters (BSR > 3) for the significant latent variable of the task PLS analysis relating $MEAN_{BOLD}$ to sample presentation.*

| | MNI | | | | | |
|---|---|---|---|---|---|---|
| **X** | **Y** | **Z** | **BSR** | **p-value** | **Cluster size (voxels)** | **Anatomical region** |
| 48 | -30 | 48 | 12.5537 | 0.0000 | 23311 | R Postcentral Gyrus |
| -15 | 51 | -15 | 7.1158 | 0.0000 | 168 | L Superior Orbital Gyrus |
| -36 (-36) | 0 (4) | -21 (-21) | 5.4933 | 0.0000 | 41 | L Temporal Pole |

*Note.* Abbreviations: MNI = Coordinates in Montreal Neurological Institute space, BSR = Bootstrap ratio. Anatomic labels were derived from the cytoarchitectonic atlas provided in the anatomy toolbox for SPM 8 (Eickhoff et al., 2005). We only report clusters of 25 voxels or more.

**Supplementary Table S6**: *Peak voxel coordinates in robust clusters (BSR > 3) for the significant latent variable of the behavioral PLS analysis relating uncertainty modulation to estimation error (N = 47).*

| MNI | | | | | | |
|---|---|---|---|---|---|---|
| X | Y | Z | BSR | p-value | Cluster size (voxels) | Anatomical region |
| 57 | -39 | 51 | 7.5835 | 0.0000 | 636 | R Inferior Parietal Lobule |
| 0 | 27 | 51 | 6.7004 | 0.0000 | 172 | L Superior Medial Gyrus |
| 48 | 39 | 15 | 6.5347 | 0.0000 | 202 | R IFG (p. Triangularis) |
| -36 | -48 | 45 | 6.3783 | 0.0000 | 365 | L Inferior Parietal Lobule |
| -42 | 39 | 15 | 6.1012 | 0.0000 | 49 | L IFG (p. Triangularis) |
| -33 | 18 | 9 | 5.8193 | 0.0000 | 40 | L Insula Lobe |
| 36 | 24 | 0 | 5.3671 | 0.0000 | 59 | R Insula Lobe |
| 51 | 12 | 33 | 5.0933 | 0.0000 | 71 | R IFG (p. Opercularis) |
| 30 | 54 | -9 | 4.5326 | 0.0000 | 27 | R Middle Orbital Gyrus |
| -48 | 6 | 36 | 4.4879 | 0.0000 | 36 | L Precentral Gyrus |
| -9 | 60 | 3 | -5.4550 | 0.0000 | 391 | L Superior Medial Gyrus |
| -18 | -6 | -21 | -4.3491 | 0.0000 | 35 | L Hippocampus |

*Note.* Abbreviations: MNI = Coordinates in Montreal Neurological Institute space, BSR = Bootstrap ratio. Anatomic labels were derived from the cytoarchitectonic atlas provided in the anatomy toolbox for SPM 8 (Eickhoff et al., 2005). We only report clusters of 25 voxels or more.

### SUPPLEMENTARY METHODS

**Task design.** During the final experimental block, performed outside the MR scanner, confidence ratings of the estimation of the blue-red marble ratio were collected. For this block of trials, a rating scale was included directly after completion of the estimation and prior to the gambling phase. Participants were instructed to indicate their confidence ranging from "very unsure" to "very sure" about the estimation of the blue-red marble ratio on a continuous confidence judgment scale [1]. Additionally, the amount of obtainable reward on each trial was manipulated. Each marble was marked with one ring (low reward) or two rings (high reward). In the high reward condition, the magnitude of the reward in the gamble was doubled compared to the low reward condition. Each participant completed each jar an equal number of times in the low and high reward conditions.

**The gambling phase.** At the end of each trial, participants had to make a risky decision between two options. One option represented a draw from the current urn. If the outcome had been a blue marble, the participant would have received a payoff with a specific magnitude (which is different in each trial, see details below). The other option represented a reference option with a certain payoff (i.e., reward probability was 100 percent). After the experiment finished, one of these gambles was picked and participants received a bonus payment based on the result of a draw from their chosen option. The options were presented in the left or right hemi-field of the screen and the order was pseudo-randomized across the experiment. Participants should choose one option via left or right button of the button box. Each trial ended following the decision and subsequently the next trial started with a different urn and reward probability.

The 18 different marble jars in the experiment differed in expected value in the following way: In six jars the expected value (probability of drawing a blue marble times the varying payoff) was larger than the reference option (ranging between 10 and 30 percent), in another six urns the expected value of both options were approximately equal and in the final six urns the expected value was lower than the reference option (ranging between -10 and -30 percent, Supplementary Table S1). The probability of drawing a blue marble varied between 0.1 and 0.91, whereas the payoff varied between 8 and 130 points. Theoretically, a decision-maker that always chooses the gambling option could expect the same summed expected value as a decision maker that always chooses the certain reference option.

### SUPPLEMENTARY RESULTS

**Model and parameter recovery.** The ground truth model was recovered well from each simulated dataset. The respective ground truth model consistently had the lowest BIC overall (supplementary Table S2). Across the simulated subjects, the ground truth model was always recovered most frequently (Supplementary Figure S1E). However, there was some model confusion, particularly for our winning model (fitted prior with noisy choice rule), which was only recovered in about 49% of simulated subjects. This may also explain the variance in empirical model fit on the subject level, although the majority of subjects (35%) were still best fit by the overall winning Bayesian observer model (Supplementary Figure S1A). Some degree of model confusion is to be expected since all variants of our Bayesian observer model aim to account for the same behavioral pattern (biased estimation of extreme marble proportions). Certain parameterizations of the prior model family may thus yield simulated behavior that is also well accounted for by models from the exponential family. Furthermore, some confusion between nested models is not necessarily surprising. For example, certain ranges of the response noise parameter $\sigma$ may result in similar behavioral predictions to that of the same model that samples responses from the final beta distribution. It is noteworthy, however, that in general models including a noisy response rule are well differentiable from models with a sampling response rule. This supports the inclusion of the noise parameter $\sigma$ in our winning model. There was also good differentiation between the Rescorla-Wagner model and all Bayesian observer models on the subject level supporting our inference that subjects seem to use uncertainty to inform their marble ratio estimates.
We also checked the recoverability of model parameters. For our winning model, both prior (Pearson's $r = 0.98$) and noise (Pearson's $r = 0.97$) parameters estimated for each simulated subject were highly correlated with their ground truth values (Supplementary Figure S1C). Parameters of the second-best model were also recoverable (Supplementary Figure S1D). This reinforces the inferences made from individuals' parameter estimates in our results.

**Explaining the effect of response noise parameter $\sigma$ on extreme jar bias**. The significant main effect of response noise $\sigma$ in predicting subjects' extreme jar bias suggests that people with opposite extreme jar biases (i.e., more misestimation for jars with marble ratios close to 50:50), is not captured by the winning behavioral model. Indeed, if we remove subjects with opposite extreme jar bias (i.e., negative effect slopes, $N = 9$) from this regression analysis, the main effect of noise parameter $\sigma$ on extreme jar bias is not significant (t(39) = -0.113, p = 0.922). Thus, our behavioral model accounts well for the empirical response pattern we sought to capture (i.e., positive effect slopes).

**Analysis of BOLD signal modulation by uncertainty**. Our PLS model relating BOLD signal modulation by uncertainty (i.e., standard GLM beta estimates for the parametric uncertainty regressor) to subjects' median estimation error revealed a significant latent association (permuted p = $3.40 \cdot 10^{-2}$, Spearman's r = 0.675, Figure 5B, cf. Figure S3B for N = 51). Because brain saliences were both positive and negative, individual differences in parametric modulation effects related differently to estimation errors between regions (see Figure S2C for full axial brain plots and Table S5 for peak voxel coordinates in robust clusters). In robust clusters with positive saliences, subjects who down-modulated BOLD activity more with increasing state uncertainty made fewer estimation errors (lower latent uncertainty modulation rank scores reflecting more negative modulation effects) than subjects who did not track state uncertainty in these regions (higher latent uncertainty modulation rank scores reflecting BOLD signal modulation by state uncertainty effects close to zero). Robust clusters were mainly located in the lateral parietal cortex, PFC (dorsomedial, ventrolateral, and rostrolateral parts), and in the insula. In contrast, in robust clusters with negative brain saliences, subjects who showed more positive coupling between BOLD signal and state uncertainty made fewer estimation errors than subjects with a negative coupling. This relationship was mainly expressed in a cluster located in the ventromedial PFC (vmPFC) and a cluster in the left hippocampus.

We also investigated whether these performance-related individual differences in BOLD signal modulation by uncertainty could be explained by idiosyncratic uncertainty representations, as captured by our winning behavioral model. We ran multiple regression predicting latent uncertainty modulation of the BOLD response, which relates to task accuracy, from the prior width parameter ρ controlling for response noise $\sigma$. This model explained a significant amount of variance (F(2,44) = 6.244, p = $4.10 \cdot 10^{-3}$) with an $R^2$ of 0.221 (Figure 5C). There was a significant main effect for prior width ρ (t(44) = 3.508, p = $1.05 \cdot 10^{-3}$, semi-partial $\eta^2$ = 0.218) but not for parameter $\sigma$ (t(44) = 1.590, p = 0.119). This finding suggests that people with suboptimal uncertainty representations, due to a narrow prior belief distribution (i.e., higher ρ), show less BOLD signal modulation by state uncertainty trajectories derived from the unbiased Bayesian observer model. We again ran this analysis also for the parameters of the (second-best) evidence weight model, which afforded similar inferences (see supplementary results).

**Brain-behavior results for the evidence weight model**. Our alternative model, which explains suboptimal responding by an exponential weighting of the incoming evidence, would similarly predict individual differences in state uncertainty reduction during the sampling phase: People who reduce uncertainty less for large samples (i.e., $\delta$ < 1) make more estimation errors than people who weight the information provided by large samples more optimally (i.e., $\delta$ = 1). Subjects' fitted prior parameter ρ of the winning model and fitted evidence weight $\delta$ for second-best model (both including a response noise parameter $\sigma$) were highly correlated (Pearson's r(49) = -0.863, p = $3.73 \cdot 10^{-16}$), supporting the notion that they capture similar behavioral response patterns. For this model, the fitted evidence weight parameter $\delta$ had a median of 0.85 (SD = 0.84) and the fitted noise parameter $\sigma$ had a median of 0.10 (SD = 0.05).

Predicting ranked latent $\Delta SD_{BOLD}$ from the ranked evidence weight $\delta$ and response noise $\sigma$ parameters in a multiple regression model explained a significant amount of variance (F(2,44) = 6.938, p = $2.41 \cdot 10^{-3}$) with an $R^2$ of 0.240. There was a significant main effect only for evidence weight $\delta$ (t(44) = -3.725, p = $5.53 \cdot 10^{-4}$, semi-partial $\eta^2$ = -0.240): People who underweight the evidence of large samples more (i.e. $\delta$ < 1) show less $SD_{BOLD}$ collapse during the sampling phase. Furthermore, the regression model predicting latent uncertainty modulation of the BOLD response from evidence weight parameter $\delta$ and response noise $\sigma$ explained a significant amount of variance (F(2,44) = 5.544, p = $7.12 \cdot 10^{-3}$) with an $R^2$ of 0.201. There was a significant main effect for evidence weight $\delta$ (t(44) = -3.257, p = $2.17 \cdot 10^{-3}$, semi-partial $\eta^2$ = -0.193) and also for parameter $\sigma$ (t(44) = 2.147, p = $3.74 \cdot 10^{-2}$). This suggests that people who underweight the evidence of large samples ($\delta$ < 1) show less BOLD signal modulation by uncertainty trajectories of the unbiased Bayesian observer.

Overall, our interpretation that individual differences in uncertainty representations explain the observed brain-behavior relationships would not have changed had we chosen this alternative model. We frame our conclusions in terms of the fitted prior model because it provided a better account of behavior.

### SUPPLEMENTARY DISCUSSION

Our standard analysis approach revealed that BOLD signal modulation by state uncertainty predicted task accuracy. On one hand, higher BOLD signal (in lateral parietal cortex, dorsomedial PFC, ventro- and rostrolateral PFC, and anterior insula) with lower uncertainty predicted higher estimation accuracy. In other words, higher performers showed an increase in BOLD activity in these regions as they observed more samples and became more certain about the jar marble ratio on a given trial. These areas correspond to frontoparietal control and dorsal attention brain networks, which support goal-directed behavior in externally-driven tasks [2-6]. These networks of brain areas have also been found to correlate with state uncertainty in other inference tasks [7,8]. However, these prior studies commonly report an opposite effect direction in which high state uncertainty reflected *more* activity in these brain regions. For example, an fMRI study by McGuire, et al. [7] investigated how uncertainty drives learning in a task that required participants to infer the position of an unseen helicopter based on bag drops that followed a normal distribution around the helicopter's true position. In their task, the helicopter location could change unannounced leading to learning rate adjustments due to change-point uncertainty and uncertainty about the helicopter location in a given environment (which the authors term relative uncertainty). The authors found BOLD signal modulation by relative uncertainty in lateral parietal cortex, dorsomedial PFC, ventrolateral PFC, and anterior insula, which match the regions we found in the current study. However, the direction of effect is reversed showing a positive coupling between relative uncertainty and the BOLD signal. Two key differences of our study are that we investigate state inference in a stable rather than dynamic environment and that decisions are only required after observing several evidence samples. In these respects, our study shares similarities with perceptual decision-making studies, which assume that noisy perceptual evidence is integrated over time to arrive at a decision [9,10]. Previous fMRI studies of perceptual decision-making also report the involvement of a similar set of higher-order brain regions, which have been linked to evidence accumulation, decision formation and response preparation [11-18]. Notably, some studies have reported an *increase* in average BOLD activity in these brain areas over the sampling period (i.e., with decreasing state uncertainty), which aligns with our findings [11,19].

Conversely, *positive* coupling between the BOLD signal and state uncertainty in vmPFC and left hippocampus also predicted better performance in our task. The previously mentioned study by McGuire, et al. [7] also reported an effect in the vmPFC, but again with opposite effect direction compared to the one we see in high performers. At first glance, our findings appear at odds with previous work reporting vmPFC tracking of subjective confidence. However, recent work by Trudel, et al. [20] suggests that representations of uncertainty in this brain region depend on the behavioral goal. In their task, participants had to learn the predictiveness of two choice options in determining a target location that was later revealed in the trial. Early on during learning, vmPFC BOLD signal positively tracked the uncertainty differences of the two choice options and correlated with uncertainty-guided exploration. Later on, vmPFC BOLD signal negatively tracked the uncertainty difference and correlated with uncertainty-avoidant exploitation. The positive effect we observe in our study is thus in line with the general idea of an "exploratory brain mode" that supports uncertainty-guided learning. Overall, the GLM results directly connect to previous studies across various domains of decision-making and reveal surprising discrepancies that require more attention in future work.
